# A Human Breast Cell Atlas Mapping the Homeostatic Cellular Shifts in the Adult Breast

**DOI:** 10.1101/2023.04.21.537845

**Authors:** Austin D. Reed, Sara Pensa, Adi Steif, Jack Stenning, Daniel J. Kunz, Peng He, Alecia-Jane Twigger, Katarzyna Kania, Rachel Barrow-McGee, Iain Goulding, Jennifer J. Gomm, Louise Jones, John C. Marioni, Walid T. Khaled

**Author notes:** Contributed equally to this manuscript (Austin D. Reed, Sara Pensa). Correspondence should be addressed to John Marioni or Walid Khaled.

## Abstract

One of the barriers for breast cancer prevention and treatment is our poor understanding of the dynamic cellular shifts that naturally occur within the breast and how these changes contribute to tumour initiation. In this study we report the use of single cell RNA sequencing (scRNAseq) to compile a Human Breast Cell Atlas (HBCA) assembled from 55 donors that had undergone reduction mammoplasties or risk reduction mammoplasties. The data from more than 800,000 cells identified 41 cell subclusters distributed across the epithelial, immune, and stromal compartments. We found that the contribution of these different clusters varied according to the natural history of the tissue. Breast cancer risk modulating factors such as age, parity, and germline mutation affected the homeostatic cellular state of the breast in different ways however, none of the changes observed were restricted to any one cell type. Remarkably, we also found that immune cells from *BRCA1/2* carriers had a distinct gene expression signature indicative of potential immune exhaustion. This suggests that immune escape mechanisms could manifest in non-cancerous tissues during very early stages of tumour initiation. Therefore, the Atlas presented here provides the research community with a rich resource that can be used as a reference for studies on the origins of breast cancer which could inform novel approaches for early detection and prevention.

## Introduction

One of the biggest challenges in treating breast cancer is the heterogeneous nature of the disease^1^. We have limited understanding of how early divergences from the homeostatic breast subtypes lead to tumour heterogeneity. Whilst large-scale cancer genomic studies indicate that different breast cancer subtypes are enriched for certain mutations^2^, not all phenotype variability and tumour behaviour can be explained by mutations alone. Studies in mice have shown that the cell-of-origin may contribute to the phenotype of the resulting tumour^3^. Defining which cell type leads to which kind of tumour is complicated by a growing list of environmental and epidemiological risk factors, such as age, which not only affect overall incidence, but also outcome^4^. Such risk factors can themselves be modulated by other factors; for example, the age-dependent risk in breast cancer is greatly reduced by pregnancy early in life^5^, whereas predisposing germline mutations (e.g. *BRCA1/2*) greatly increases age-associated risk^6^. How these risk factors interact, and impact tissue homeostasis remains to be fully understood. To do this, it is crucial to characterise cell types and cellular states present under different physiological conditions.

To tackle this problem, studies in the mouse have leveraged single cell genomics (i.e. scRNA-seq) to determine the gene expression profile of individual mammary epithelial cells across embryonic and adult developmental stages^7–10^. Similar to the mouse, after birth the human mammary gland continues to develop. We know that at birth the human breast consists of a ductal structure with well-defined terminal ductal lobular units (TDLUs) similar to what is found in the adult^11^. Prior to puberty, the breast grows in proportion to the rest of the body. The onset of puberty triggers the expansion of the TDLU to form adult lobules and to fill the surrounding stroma. Consisting of the vasculature system, fibroblasts and immune cells, the stromal compartment remains severely understudied^13^. Despite this, mounting evidence suggests that changes in the microenvironment are a major contributing factor in tumorigenesis and an important target for novel therapeutics^14^. Together, these motivate the study of the full range of complex interactions between epithelial cells and the stromal surroundings.

Recent single cell transcriptomic and proteomic studies of human reduction mammoplasty and tumour samples have begun to catalogue the various epithelial compartments in the breast^12–17^. However, larger sample sizes are needed to generate a comprehensive transcriptomic map of all adult breast cell subtypes and states, while taking into account the major developmental changes and risk-modulating factors. To this end, in this study we report the use of scRNAseq to compile a comprehensive Human Breast Cell Atlas (HBCA) of over 800,000 cells collected from 55 women across various adult development stages.

## Results

### Identification of a suitable tissue cohort

Here we present a comprehensive human breast cell atlas, mapping shifts in cellular composition associated with various biological and environmental changes. To assist further study into a wide range of these factors we identified a cohort of healthy breast tissue samples from the Breast Cancer Now Tissue bank that had (up to 80) health and lifestyle records available (**Fig 1 and Supplementary Table 1**). Our cohort consisted of tissue samples from 22 women who had undergone reduction mammoplasties, 27 women carrying a *BRCA1* or *BRCA2* mutation or other family histories (who had risk reduction prophylactic mastectomies) and 6 contralateral mastectomies from BRCA1 carriers that had breast cancer in one breast and had the second breast removed to reduce the risk of further tumours (**Fig 1**). The samples had a wide distribution of values across the various risk modifiers such as age, parity status and menopause. Additionally, we collected a range of metadata on further risk-modifying factors that may prove beneficial to future studies exploring breast cancer risk (**Supplementary Figure 1 and Supplementary Table 1**). All samples were pre-processed by the tissue bank to isolate the epithelial and stromal/immune enriched compartments (**Fig 1**). These were then processed to single cell level and viable cells were loaded on a 10x chromium chip for single cell capturing to enable scRNA-seq. For the epithelial enriched compartment, we used fluorescence-activated cell sorting (FACS), as previously described^18^, to enrich for the luminal progenitor compartment (EPCAM^+^, CD49f^+^) (**Supplementary Figure 2-4**), which have been proposed to be the cell of origin for some breast cancers^3, 18^.

**Figure 1.**
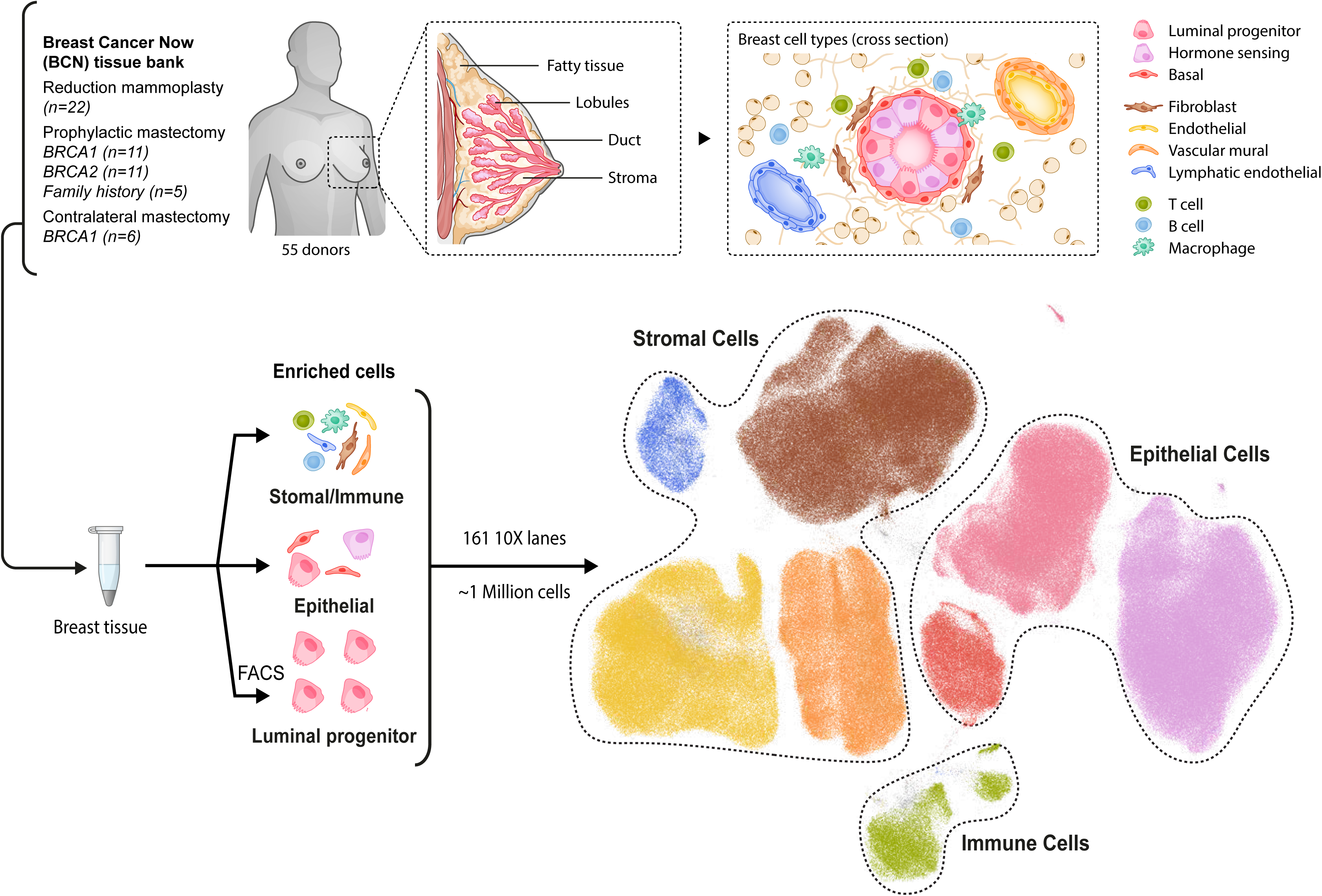
Overview of the Human Breast Cell Atlas. A schematic highlighting the overall experimental design and the cell types we aimed to capture in the Atlas. The diagram highlights the overall number of donors sequences and how they are distributed among the various subgroups. The global uniform manifold approximation and projection (UMAP) representation of the final dataset coloured by general cell types captured from all 161 samples processed as part of this Atlas.

Following sequencing, more than 1 million cells were identified and then after several quality control and doublet calling steps, 801,360 cells were taken forward for downstream analysis (**Fig 1 and Supplementary Fig 5-6**). To combine our 45 separate batches of sequencing we used scVI^19^ to produce a batch corrected embedding for use in downstream analysis. More details on the sample preparation and analysis can be found in the material and methods section. Coarse cell type annotation revealed that we sequenced over 350,000 epithelial cells, and 420,000 stromal and immune cells (**Fig 1**).

### Major cellular groups identified in the Human Breast Cell Atlas

Within the epithelial compartment we identified the Luminal Progenitor (LP), Hormone Sensing (HS), and Basal (BSL) cell types using canonical lineage markers (**Fig 2a,b**). Iterative clustering identified several subclusters within each cell type based on unique gene expression profiles. The majority of these subclusters were found in all 55 donors, albeit to varying proportions (**Fig 2c, Supplementary Fig 7a, d-h**). Of note, we found no distinct LP cluster strongly marked by alveolar-like genes (*CSN2/3*, *LALBA*; **Supplementary Fig 8d**). Instead, we find most LP heterogeneity is distinguished by proportions of standard marker gene expression. LP1/2/3 show high expression of LP specific markers (*ALDH1A3, SLPI*) in contrast to LP4. Additionally, LP2 is notable for its co-expression of both LP (*ALDH1A3*) and BSL markers (*KRT14, KRT5*) (**Supplementary 8a, d**). We also identified LP5 as a small population of proliferating LP cells marked by the canonical proliferation marker *MKI67* and mitosis related genes (*AURKB, TOP2A*; **Supplementary Fig 8a, d**). Similarly, within the HS compartment, upregulation of *AREG* marked HS1 cells, upregulation of estrogen and progesterone receptors (*ESR1* and *PGR*) marked HS2 cells, whilst high *ANKRD30A* expression marked HS4. Subcluster HS3 shows an expression pattern distinguished by high *SERPINA1 and PIP* expression alongside increased expression of some LP markers (*ALDH1A3*, *SLPI*; **Supplementary Fig 8b, d**). A further two Donor Derived Clusters (DDC) were identified, each consisting primarily of cells from just one donor (Donor 37 and Donor 17 respectively). Both DDC1 and DDC2 show high expression of LP marker genes (*ALDH1A3* and *SLPI* but not *KIT*). Despite this, they also show a mix of other epithelial lineage markers from HS and BSL cell groups with DDC1 showing particularly high expression of *MMP3* alongside *PIP* and *MUCL1*, which were otherwise predominantly expressed in HS3 (**Supplementary Fig 8d**). Despite the expression of mixed lineage markers, both DDC1 and DDC2 were not associated with batch, have low doublet scores and typical quality control metric distributions **(Supplementary Fig 8e-h**). This suggests that they represent a genuine and distinct donor specific clusters which might be indicative of early signs of disease.

**Figure 2.**
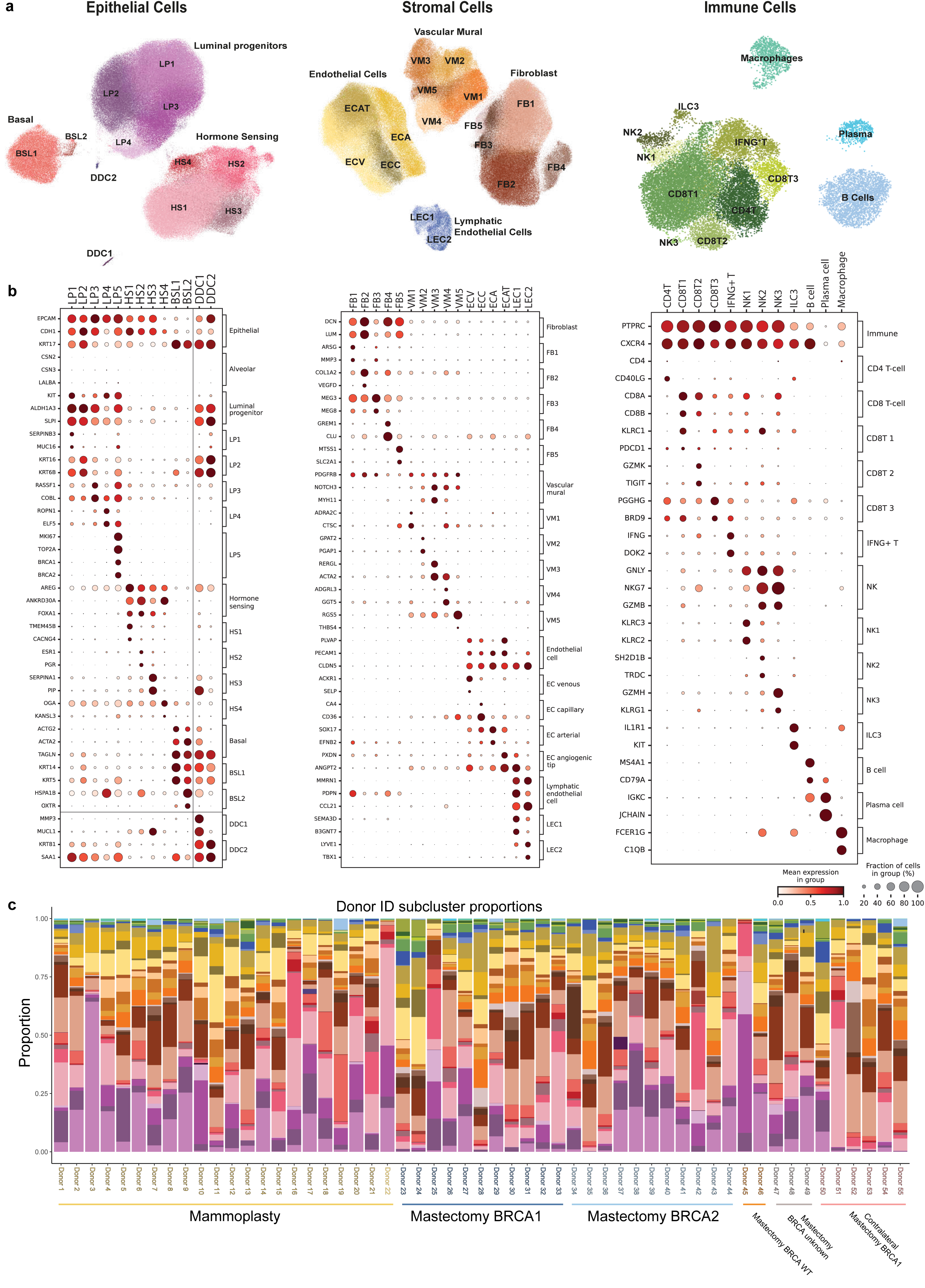
Major cellular groups identified in the Human Breast Cell Atlas. **(a)** Uniform manifold approximation and projection (UMAP) plots of the epithelial, stromal and immune cell compartments coloured and labelled by subcluster annotations. **(b)** Dot plots summarising the known markers used to identify cell types and a selection of genes distinguishing each subcluster for the epithelial (left), stromal (middle) and immune (right) compartments. Each row corresponds to a specific cell subcluster and its expression of each gene (column) normalised per gene, brackets on the right side each plot detail the cell type/subcluster that this gene marks. **(c)** Stacked bar plots displaying the proportional cellular composition of each donor normalised on a per donor basis.

Within the stromal compartment we identified five Fibroblast (FB), four vascular Endothelial Cell (EC), two Lymphatic Endothelial Cell (LEC) and five Vascular Mural (VM) subclusters present across all donors (**Fig 2a-c and Supplementary Fig 7b, d-h**). FB1 is distinguished by expression of various genes related to extracellular matrix (ECM) disassembly (*MMP2*, *MMP3, MMP10 MMP12* and *SH3PXD2B*), whereas FB2 and FB4 show increased expression of genes related to ECM formation (*DCN*, *LUM*, *COL1A1/2*; **Supplementary Fig 9a, d**). The venous, arterial, capillary and angiogenic tip EC subclusters were identified using a range of previously described marker genes^20–22^. The LEC compartment, distinguished by canonical markers *CCL21* and *PDPN*, splits into two groups that can be separated by their expression of chemokines (*CXCL1* and *CXCL8*) and some angiogenic tip cell markers (*ANGPT2*, *PXDN*) both of which are highly expressed in LEC1, but lowly expressed in LEC2 **(Supplementary Fig 9b, d**). The VM subclusters appear in two main groups with VM1/2 having low expression of genes relating to muscular functions (*ACTA2*, *TAGLN2*) and pericyte marker (*RGS5*)^23^ which are high in VM3/4/5 **(Supplementary Fig 9c, d**). Using previously described immune cell markers we were also able to identify a variety of lymphoid and myeloid cell types including five T cell subsets, three NK cell subsets, ILC3 cells, B cells, plasma cells and Macrophages (**Fig 2a, b and Supplementary Fig 10a, b**). To provide robust 100-gene signatures for each identified subcluster, we used pseudo-bulk one-versus-all differential expression testing to identify an extensive list of marker genes. These lists show high specificity to our cell type subclustering, offering many potentially novel cell type markers **(Supplementary Fig 11a and Supplementary Table 2**). We found all major cell types represented in all donors regardless of BRCA, age or parity status (**Fig 2c and Supplementary Fig 7**).

### Changes in healthy breast composition under different physiological conditions

We first consider how natural factors such as age and parity impact the composition of the breast. To avoid confounding our results with changes resulting from differences in BRCA status, we exclusively consider the 22 reduction mammoplasty donors. To gain high-resolution insight into cellular differential abundance and compositional shifts we used Milo’s flexible generalised linear models, thereby avoiding both statistical confounding and the inherent biases placed by fine cell type clustering^24^. The majority of the significant changes driven by age occur within the subclusters of the epithelial compartment. In the LP compartment, older samples display an enrichment of LP2/4 cells, while regions of the LP1/3 subclusters are enriched in younger individuals (**Fig 3a, b).** The LP clusters enriched in older women have reduced expression of traditional LP markers, consistent with previous studies showing increased LP lineage infidelity with age^14, 25^. In the HS compartment, we found an enrichment in the HS2/4 and to a lesser extent HS3 subclusters with age while in the BSL compartment we found a significant decrease in the proportion of BSL1 cells (**Fig 3a, b).** Outside the epithelial compartment, there were very few significant differences except for an enrichment in FB4 and a slight decrease in the proportion of EC angiogenic tip cells (**Fig 3a, b**).

**Figure 3.**
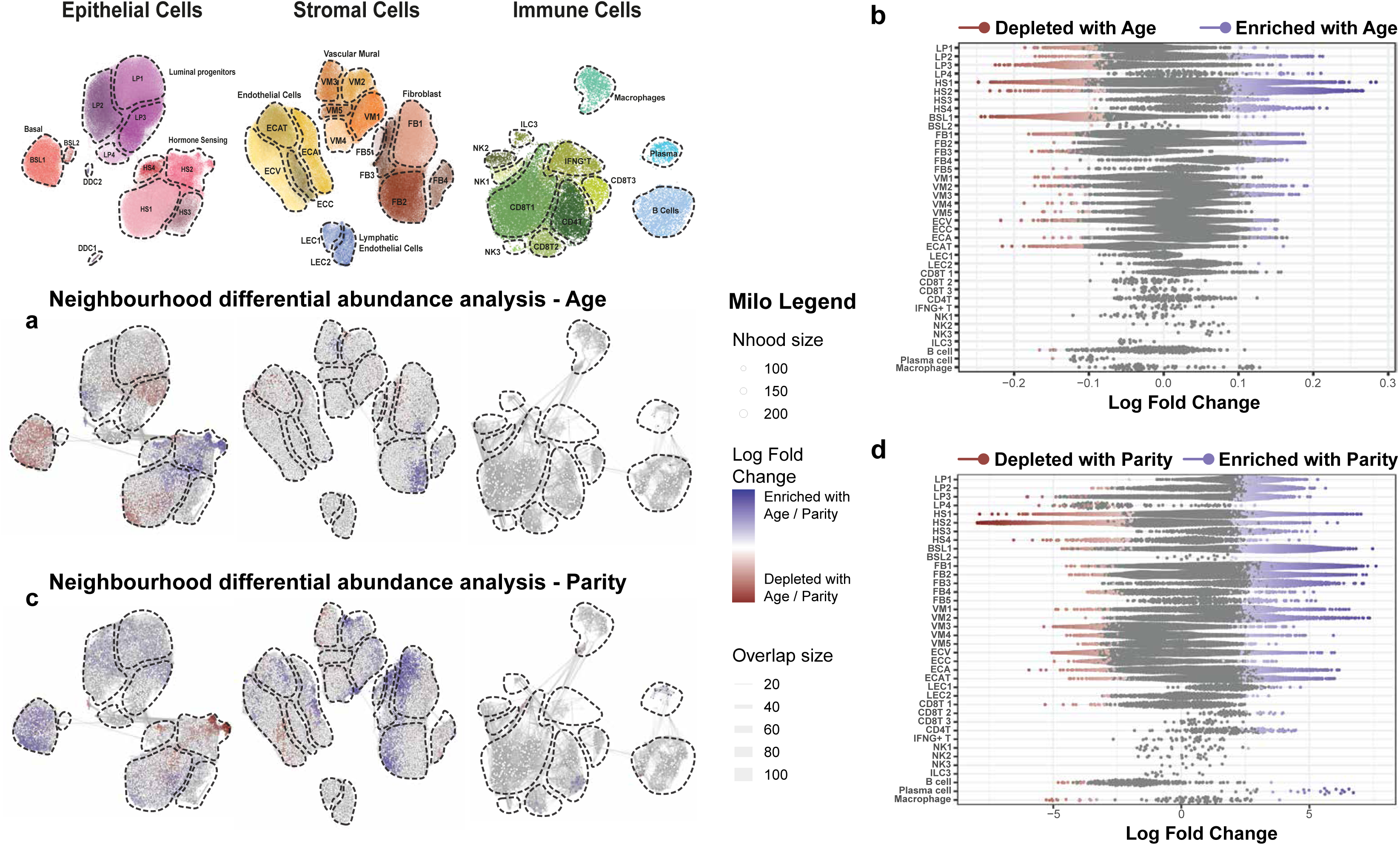
Age and parity affect the homeostatic cellular state of the breast. **(a)** Milo cell neighbourhood differential abundance plots of the significant (FDR < 0.05) changes in the breast composition with age (mammoplasty donors; n=22), blocking for the effects of parity. We test the effects of age as a continuous scale ranging from 19-65 years, with the colour gradient scale representing log fold changes per year. Blue represents enrichment with age whilst red denotes depletion with age. **(b)** Beeswarm plot of the log fold changes in the Milo neighbourhoods with age, grouped into each cell type subcluster. Neighbourhoods with a significant change in cellular abundance are coloured as indicated. Log fold changes are per year due to the continuous age scale tested. **(c)** Milo cellular neighbourhood differential abundance plots of the significant (FDR < 0.05) changes in the breast composition with age (mammoplasty donors; n=22), blocking for the effects of parity. Blue represents enrichment with parity whilst red denotes depletion with parity. **(d)** Beeswarm plot of the log fold changes of the Milo neighbourhoods with parity (*i.e*. nulliparous versus parous), grouped into each cell type subcluster. Neighbourhoods with a significant change in cellular abundance are coloured as indicated.

In contrast, parity has a more widespread impact on the cellular composition of the epithelia, stromal and immune compartments (**Fig 3c, d**) which could reflect the large-scale tissue remodelling of the breast that occurs during pregnancy. In the LP compartment, there is a decreased proportion of LP4, despite enrichment for the LP1/2/3 subclusters in women with children. In the HS and BSL compartments, there are enrichments for the HS3 and BSL1 subclusters respectively. We note that many of the changes in enrichment seen within the epithelial compartment are in the opposite direction when considering age or parity as the covariate, which could contribute to contrasting breast cancer risk posed by each. We see some evidence of this in overall increased proportions of plasma, CD8T2/3 and CD4T cell types in the profiled parous donors. In the stromal compartment, our findings suggest a transition from *ACTA2* expressing VM3/4/5 towards VM1/2-type vascular mural cells as a function of parity, as well as increases in EC arterial / EC angiogenic tip cells and an overall increase in all FB subclusters excluding FB4 (**Fig 3c, d**).

### Impact of high-risk BRCA1 and BRCA2 germline mutations on breast cellular composition and transcriptional profile

To determine mammary cell state shifts in high risk (HR) donors compared with average risk (AR) donors, we analysed cell profiles from reduction mammoplasty donors (n=22) and donors with *BRCA1 (n=11)* and *BRCA2 (n=11)* germline mutations (denoted HR-BR1 and HR-BR2 respectively). First, we test for differential expression between AR and HR-BR1 as well as between AR and HR-BR2 cohorts, accounting for the effects of both age and parity (see methods for details). We observe 429 (LP), 141 (HS) and 100 (BSL) genes upregulated in the respective HR-BR1 cell types, compared to 61 (LP), 41 (HS) and 0 (BSL) upregulated genes in the HR-BR2 donors (FDR < 0.05 and logFC > 1) (**Supplementary Table 3**). Despite similar statistical power, we noted that HR-BR1 epithelial cells have more significant transcriptional changes than the respective HR-BR2 cells. This is particularly clear in the BSL compartment which showed no significantly upregulated genes in HR-BR2 cells compared to the AR cells.

Relative to AR controls, we note significant increases in programmed cell death 1 ligand 1 (*PDL1* or *CD274*) in HR-BR1/2 LPs and HR-BR1 HS cells (**Fig 4b, Supplementary Fig 12a, b, d and Supplementary Table 3**). However, *PDL1* expression is rare, occurring on average in less than 5% of each epithelial subtype **(Supplementary Fig 12k-m**). Similarly, despite its rare expression, we also see evidence for the upregulation of milk-biosynthesis or alveolar-like genes (*CSN2/3, CSN1S1* and *LALBA*) in both HR-BR1 and HR-BR2 LP cells **(Supplementary Fig 12a, d**). The upregulation of milk-biosynthesis genes is similarly interesting to tumour development, with reference to recent mouse studies proposing these genes as a possible markers of pre-tumour progression^10, 26^. This data suggests a similar process may also characterise early signs of malignant progression in the human breast. Unsurprisingly, the expression of milk-biosynthesis genes also appears to have a strong relationship to parity, with a strong upregulation in parous donors regardless of BRCA status **(Supplementary Fig 12g, h, I, j)**.

**Figure 4.**
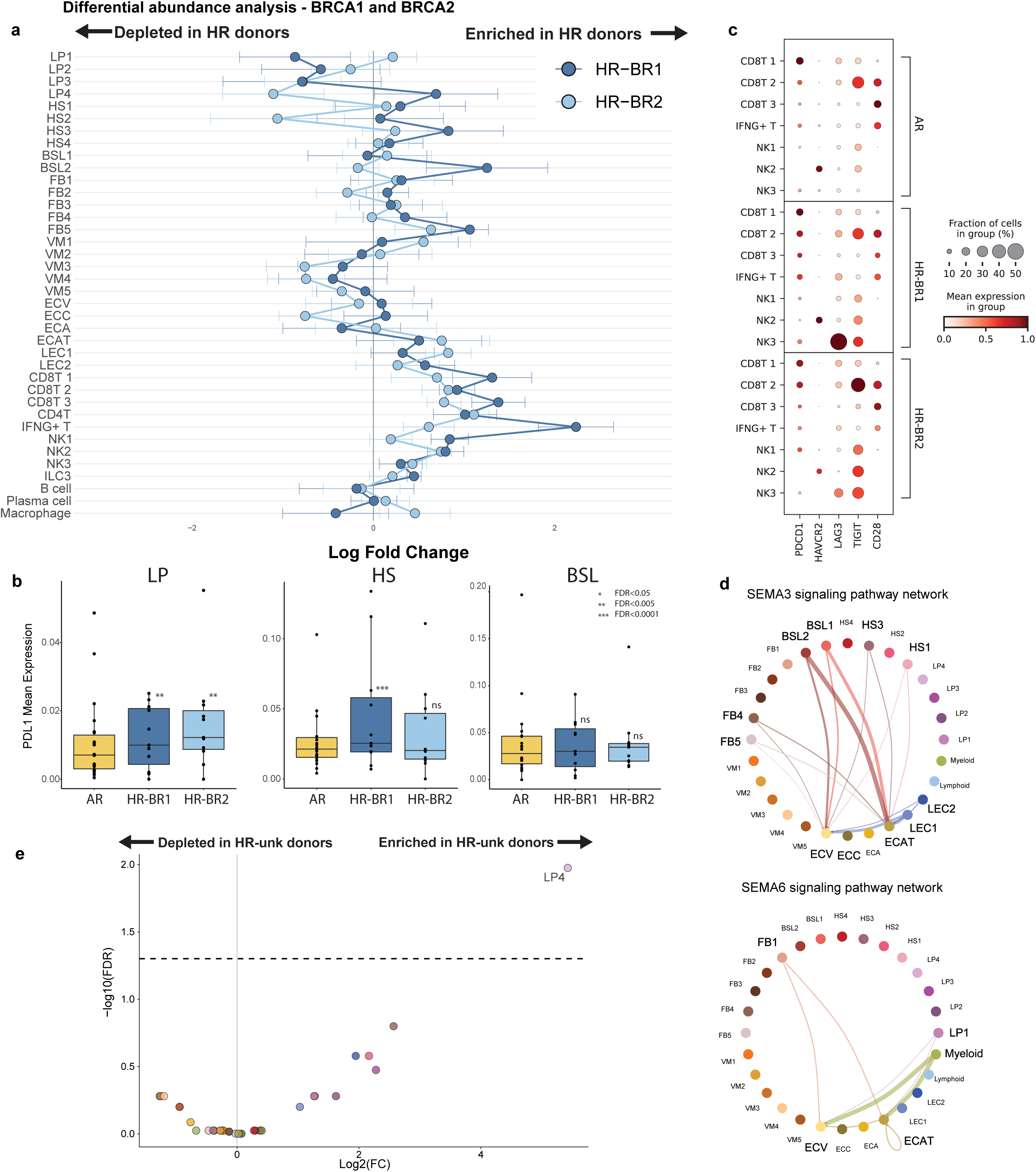
Impact of high-risk *BRCA1 (HR-BR1)* and *BRCA2 (HR-BR2)* germline mutations and high-risk (HR) germline predispositions of unknown origin on breast cellular composition and transcriptional profile. **(a)** Summary plot of average Milo differential abundance results comparing the cellular composition of the breast from average risk (AR) donors against HR-BR1 and HR-BR2 donors, blocking for the effects of both age and parity. To enable fair comparisons, neighbourhoods were computed to be shared by AR/HR-BR1 and AR/HR-BR2 tests. The plot summarises the mean and variance of log fold changes for each cell subcluster in both the HR-BR1 and HR-BR2 cohorts. A zero log fold change represents the same subcluster proportions as AR donor tissue on average, whilst positive (negative) log fold changes denote subcluster enrichment (depletion) in the relative HR donor breast. See **Supplementary Figure 15** for full Milo results in HR comparisons. **(b)** Boxplots visualising the change in mean expression (log-transformed counts) for immune checkpoint ligand *PDL1*, identified as significantly upregulated in HR-BR1 (LP and HS) and HR-BR2 (LP) donors. The FDR values are derived from negative-binomial differential gene expression testing results (see methods and **Supplementary Figure 1**2). **(c)** Dot plot displaying the expression of several key immune checkpoint receptors expression in a range of cytotoxic lymphoid subclusters comparing between AR, HR-BR1 and HR-BR2 donor groups. Expression is normalised per gene. **(d)** Cell Chat circle plots displaying the predicted ligand-receptor interactions of SEMA3 and SEMA6 signalling pathways in HR-BR2 donors. These were selected as the two most specific incoming cell-cell interaction pathways to the EC angiogenic tip cells (see Supplementary Fig 16). **(e)** Differential abundance volcano plot comparing AR to high-risk unknown germline predisposition (HR-unk) donors (blocking for age and parity) showing the difference in relative abundance of each subcluster. The points are coloured by their subcluster and identified as significant when FDR < 0.05 and the log fold change is greater than 1.

To explore other cellular differences particularly in the immune and stromal compartment, we use Milo to compare average-risk (AR) reduction mammoplasty against HR-BR1 or HR-BR2 donors and identify compositional changes that may contribute to the higher risk of breast cancer development. To allow easy comparisons between HR-BR1 and HR-BR2 induced changes we summarised this in a ‘Milo signature’ plot averaging neighbourhood log fold changes per donor for each subcluster (**Fig 4a and Supplementary Fig 15**). In the HR-BR1 cohort, one of the most significant changes is the large increase in proportions of immune cell subclusters CD4 T cell, CD8 T cell 1/2/3 and NK1/2 (**Fig 4a**). We also find a two-fold enrichment in a *CD8A/B*^+^ cell type distinguished by high *IFNG* expression, suggesting a CD8 T-cell phenotype with a potential pro-inflammatory role (**Fig 4a and Supplementary Fig 10a, b**). Gene set enrichment analysis (GSEA; see methods) of the top 100 differentially expressed genes defining this population (**Supplementary Table 2**) reveals “immune response”, “T-cell activation”, “T-cell differentiation” and “positive regulation of cell killing” as some of the most significantly enriched Gene Ontology biological processes **(Supplementary Fig 13a**). This supports the hypothesis of a pro-inflammatory role and suggests that the expansion of this population of T cells is likely driven by a direct immune response.

To look for the possible cause and impacts of this immune response we turned towards the epithelial compartment. There is evidence to suggest population shifts in each of the HS cell subclusters, with a significant increase in the HS3 population. The strongest effects are seen in the increase in proportions of LP4 and BSL2 cell types, paired with relative decreases in the remaining LP and BSL subclusters (**Fig 4a and Supplementary Fig 15**). Looking at the top 100 differentially upregulated genes in LP4 vs all other LP cells revealed a range of genes relating to stress and heat shock response (*JUN*, *FOS*, *HSPA1A*) alongside *RECQL*, *FBXW7*, *MCL1* and *GADD45B*/G **(Supplementary Fig 13b, Supplementary Table 4**). *RECQL* and *FBXW7* are putative breast cancer tumour suppressors that play a role in maintaining genomic stability and the DNA damage response^27, 28^. *GADD45B/G* on the other hand are known to be induced in response to DNA damage^29^ and *MCL1* is involved in the recruitment of the double strand repair machinery^30^. A similar expression profile is seen in the BSL2 and HS2 populations **(Supplementary Fig 13b, Supplementary Table 4**) suggesting that each of these populations may be harbouring DNA damage. Given BRCA1’s instrumental role in maintaining genomic stability, this could provide a possible link between the donor HR-BR1 status and these compositional changes.

With the aim of better understanding observed epithelial cell changes in HR-BR1 donors, we performed single-cell regulatory network inference and clustering (SCENIC)^31^ to identify unique transcription factor (TF) networks regulating each subcluster **(Supplementary Fig 14a-c**). We found several regulons that were highly expressed in each of the LP subclusters including *CEBPD*, *STAT1* for LP1, *GLIS3*, *FOXJ3* for LP2 and *TEAD1*, *CEBPG* for LP3 **(Supplementary Fig 14a, d, e)**. Subcluster LP4 displayed a distinct regulon expression pattern, with regulation of transcription factors such as ISL1 and ETV1, which have been shown to have an oncogenic role in breast cancer^32, 33^. Additionally, LP4 shares several highly expressed transcription factor regulons with the LP5 cell subcluster including BRCA1, MYBL1, EZH2 and E2F2/7/8 **(Supplementary Fig 14a, e**), all of which are known to play important roles in the DNA damage response, cell cycle and breast cancer^34–36^. In particular, EZH2 expression is known to have specific links to *BRCA1* driven breast cancers and has been reported to increase the efficacy of double-strand-break-inducing platinum-based chemotherapies in mice^37^. The BSL and HS compartments show concordance with known transcription factor cell type markers with high expression of TP63^38^ and FOXA1^39^ regulons, respectively (**Supplementary Figure 14b, c, f, g**). Together these findings identify a range of novel epithelial subcluster specific regulons which may be used in the future to uncover the regulation and function of these subtypes **(Supplementary Fig 14**).

Similar to the HR-BR1 cohort, HR-BR2 donor cells display an increased immune response with the expansion of CD8T 2/3, CD4T and IFNG+ T (but not NK1/2) cell types, albeit not to the same extent as observed in HR-BR1 donors (**Fig 4a and Supplementary Fig 15**). The expansion of IFNG+ T cells seen both in HR-BR1 and HR-BR2 donors could thus correlate with the previously mentioned upregulation of *PDL1* in epithelial cells (**Fig 4b and Supplementary Fig 12a, b, d**). To look for signs of increased cytotoxic lymphoid cell exhaustion and inactivation we looked at the difference in expression of a range of immune checkpoint genes^40^ including *PDCD1*, *HAVCR2*, *LAG3*, *TIGIT* and *CD28*. We see general increases in the frequency and extent of expression across our immune checkpoint genes in both HR-BR1 and HR-BR2 donors (**Fig 4c**). There are modest increases in the expression of *PDCD1* (PDL1 receptor) itself in the CD8T cell 2, IFNG+ T (HR-BR1 specific) and NK1 subclusters, alongside increased expression of *LAG3* and *TIGIT* in the CD8 T and NK cell types of HR-BR1/2 donors (**Fig 4c**). This offers further evidence that whilst there is increased immune response in the HR donors, the immune system is showing signs of exhaustion and suppressed function, in these precancerous tissues.

Contrasting to the results found in the HR-BR1 cells, the HR-BR2 donors show evidence of a decrease in both the LP4 and HS2 subclusters. Perhaps surprisingly, the most significant changes observed in the HR-BR2 cohort appear in the stromal compartment. Here, we see significant shifts within the VM compartment from VM3/4/5 towards VM1/2, mirroring some of the changes seen with parity, alongside an enrichment of FB5 cells (**Fig 4a and Supplementary Fig 15**). However, the strongest effects are seen amongst the two endothelial compartments with shifts towards the EC angiogenic tip and LEC1 populations (**Fig 4a and Supplementary Fig 15**) suggesting dysregulated vasculature and lymphatic development. Excluding the HR-BR2 Donor 37, which harbours the outlier population DDC1, all other donors show no evidence of aberrant or malignant epithelial lesions. Interestingly, this suggests that our HR-BR2 donors have a perturbed vascular microenvironment seemingly predating aberrant epithelial expansion.

To explore the possible contribution of cell-cell communication in driving the enrichment of angiogenic tip cells we used Cell Chat^41^ to analyse the co-expression of known ligand-receptor pairs indicating putative cell-cell interactions. First, comparing global changes between AR and HR we have summarised the difference in overall incoming and outgoing signalling from each subcluster **(Supplementary Fig 16a**). The most notable changes shared by the AR vs HR-BR1 and AR vs HR-BR2 comparisons are large decreases in incoming signalling to the HS1/3 subclusters and increased incoming signalling to LP1/3 cells, with the latter effect being much stronger in HR-BR1 donors. A general increase in FB1 incoming and outgoing signalling was also observed. There are few overall signalling changes specific to the HR-BR2 comparison excluding a slight increase in outgoing VM1/2 signalling **(Supplementary Fig 16a**). Thus, we opted to further investigate individual pathways communicating with the EC angiogenic tip cells.

We identified the top 12 incoming signalling pathways specific to the EC angiogenic tip cells revealing a range of pathways well known to be involved in angiogenesis such as VEGF, ANGPT and VISFATIN whose expression appears relatively ubiquitous and difficult to trace **(Supplementary Fig 16b, c**). Two of these pathways, SEMA3 and SEMA6, were found to be highly specific to the EC angiogenic tip cells **(Supplementary Fig 16b**). Using circle plots we see that SEMA3 signalling is driven by BSL1/2 cells while SEMA6 signalling is driven by the FB1 and myeloid subclusters (**Fig 4d**). Despite originally being known for their role in neuronal development, recent studies have shown the importance of semaphorins in vascular maintenance and development. SEMA3, particularly SEMA3A, is thought to have a role in anti-angiogenesis and vascular normalisation^42, 43^ while SEMA6 is reported to be involved in driving adult angiogenesis^44^. Of these, Cell Chat information flow testing showed a significant increase in SEMA6 and ANGPT signalling pathways into the EC angiogenic tip cells, alongside a decrease in overall VEGF signalling **(Supplementary Fig 16d**). Together this suggests that aberrant SEMA6 and ANGPT signalling could be contributing to the observed vascular abnormalities in HR donors.

To explore how changes in the HR-BR1/2 cohorts might compare to other HR backgrounds we also considered the compositional changes in our donors with breast cancer predispositions of unknown origin (denoted HR-unk; including two donors testing WT for both BRCA1 and BRCA2). To ensure we could reliably block for the effects of age and parity with lower sample numbers, we were unable to run high-resolution Milo testing and instead opted for a more traditional negative binomial general linear model using edgeR. This analysis revealed that only the LP4 subcluster to be significantly enriched (**Fig 4e**). Interestingly, similar LP4 enrichment was also seen in our HR-BR1 but not HR-BR2 cohorts. This suggests that this population might play an important role in mediating or as a direct consequence of the increased risk of tumour formation across donors with different genetic predispositions to disease.

## Discussion

In this study we describe a scRNAseq Human Breast Cell Atlas generated by sequencing over 800,000 cells from 55 donors. The scale of this dataset has enabled us to delve into the entire breast composition, encompassing not only the epithelium but also the surrounding microenvironment. Due to the diversity of the samples, we were able to interrogate the data in relation to several key breast cancer risk modifiers, such as age, parity and *BRCA1/2* germline mutations, enabling us to uncover cellular interactions and compositional changes associated with each factor.

As expected, both age and parity affect the homeostatic cellular state of the breast. Although the changes observed are not restricted to any one type of cell, we could identify unique features for the two risk factors. In contrast to parity, which has a widespread impact on breast composition, the main changes associated with age are concentrated within the epithelial compartment. Most notably, we observe an enrichment of the LP2/4 clusters, which are characterised by mixed lineage expression and reduced loyalty towards traditional LP markers. This is in line with previous observations of increased LP lineage infidelity with age^14, 25, 45, 46^. The global impact of parity can be seen across the epithelial, stromal and immune compartments. Many of the effects observed in the epithelium oppose those observed with age such as strong changes in BSL1 and HS2 proportions alongside more subtle general shifts in the total proportion of LP cells. Outside the epithelium, we note increased proportions of plasma, CD8T2/3 and CD4T cells in parous women (**Fig 3**). Overall, we found that the cellular composition changes observed with parity and age are complex and not restricted to any one type of cell. Thus, the impact of these changes needs to be considered collectively rather than in isolation given the contrasting impact of age and parity on breast cancer risk^4, 47^.

Once age and parity are accounted for, we do not identify cell populations that are exclusively associated with *BRCA1/2* germline mutation carriers, rather we observe shifts in the proportions of certain cell type subclusters. One of the most diverse cell populations we captured in this study is the luminal progenitor compartment which is also the sub population of epithelial cells most associated with breast cancer^18^. Some of the changes we observed here include decreased overall LP proportions in both HR groups, with *BRCA1* samples having an enrichment of the LP4 population, similar to that seen with age. Additionally, there was strong enrichment for the HS3 and BSL2 subclusters in HR-BR1 donors that was not seen in the HR-BR2 donors (**Fig 4a and Supplementary Fig 15**). Some of the largest changes seen in our HR cohorts occurred in the immune compartment. Particularly, we detected a significant immune expansion, prominently of the IFNG+ T cells in HR-BR1 donors, accompanied by increased expression of canonical immune checkpoint receptors (ICRs) in both HR cohorts (**Fig 4c**). Interestingly, when considering wide scale transcriptional shifts in the epithelial cells of the HR donors compared to AR, we also observe a slight increase in *PDL1* expression, mainly in LPs but also across some of the HS cells (**Figure 4b and Supplementary Figure 12**). Previous studies have shown that high expression of *IFNG* in CD8 T cells, resembling what we observe in the IFNG+ T cell population, can directly drive *PDL1* expression in melanoma^48^, suggesting that this expansion of IFNG+ T cells could be contributing to the *PDL1* induction in HR-BR1 donors. The upregulation of *PDL1* could be particularly interesting due to its key role in immune evasion and supporting a tumourigenic microenvironment^49^. Given the observed increase in ICRs in HR donor immune cells and the accumulating evidence of the involvement of LPs in tumour initiation, our findings point towards an early epithelial immune-escape mechanism in the pre-malignant tissue driven by *BRCA1* and *BRCA2* germline mutations.

Within the stromal compartment, we found a strong enrichment in EC angiogenic tip cells in HR donors, particularly HR-BR2 (**Fig 4a and Supplementary Fig 15**). Cell-cell interaction analysis suggests that SEMA3 and SEMA6 signalling pathways could be involved in mediating this enrichment (**Fig 4d**). This agrees with a previous study where male *BRCA* mutation carriers were found to have increased proportions of endothelial progenitor cells^50^. This warrants future investigations given the role of angiogenesis and vascular remodelling in tumourigenesis.

Finally, one of our primary objectives was to provide the community with a robust reference dataset that would enable seamless projection and integration with other datasets (materials and methods). As an example of this potential, we have integrated the scRNAseq dataset obtained from cells isolated from mammoplasty donors and human milk^15^, finding that the common cell types cluster as expected **(Supplementary Fig 17**) demonstrating the potential for expanding the atlas as more datasets become available. In summary, we have generated a rich scRNAseq atlas of the adult human breast which has revealed novel insights into the cellular changes associated with age, parity and germline mutations as well as corroborating results from other single cell genomic studies^12, 14–17^.

## Methods

### Human tissues

All primary human breast tissue was obtained from the Breast Cancer Now Tissue bank (REC 15/EE/0192).

### Batch design

Samples were divided into 45 batches (labelled by processing_date in adata/sce object) where each batch represents a day in which 2 (for epithelial-enriched fractions) or 4 (for stromal-enriched fractions) samples were processed together. The batches were designed to minimise confounding with any of the main demographics of age and parity, or tissue conditions (surgery type or BRCA status). To achieve this, batches were designed to be either age or parity matched but with the corresponding tissue condition pseudo-randomised.

### Mammary gland dissociation into single-cell suspension

Frozen vials of epithelial-enriched or stromal-enriched fractions were defrosted by gently diluting the material in 50 mL of cold Tissue Preparation Medium (TPM, RPMI 1640 + 25 mM HEPES and 2 mM L-glutamine (Sigma R5886), 5% foetal bovine serum (FBS, Gibco), 100 units/mL penicillin and 0,1 mg/mL streptomycin sulphate (Gibco), washed in PBS without Calcium and Magnesium (D8537, Sigma). Samples were centrifuged at 400 g for 5 minutes and resuspended in 2 mL of freshly prepared PBS + 0,025% Trypsin, 0,1 g/L EDTA (HyClone SV30031.01, Fisher Scientific) and 0,4 mg/mL Deoxyribonuclease 1 (DNase) (10104159001, Boehringer/Roche Diagnostics) previously warmed to 37°C. Samples were then incubated at 37°C with pipetting up and down for 30 seconds every 2-3 minutes until smoothly digested or up to a maximum of 10 minutes. Next, samples were washed in 40 mL of TPM and centrifuged for 20 minutes at 400 g with slow break. The pellet was resuspended by pipetting up and down in 200 μL of TPM + 10 μL of 10 mg/mL DNase until homogeneous, then diluted in 25 mL of TPM and filtered through a 40 μm cell strainer (352354, Corning) into a 50 mL tube. After centrifugation for 5 minutes at 400 g, the pellet was resuspended by pipetting up and down in 200 μL of CPM (Cell Preparation Medium, RPMI 1640 + 1% FBS, 100 units/mL penicillin, 0,1 mg/mL streptomycin sulphate) + 10 μL of 10 mg/mL DNase until homogeneous, then washed in 3-6 mL of CPM. 30,000 cells were resuspended into 48 μL of HF (Hank’s balanced salt solution (Gibco) + 1% FBS) into low binding tubes. 400 Human mammary epithelial cells (HuMECs, Thermo Fisher Scientific A10565) were added as spike-in, and samples were submitted for scRNAseq (unsorted fraction). For the epithelial-enriched fraction only, the rest of the processed sample was stained with the following primary antibodies: CD45-APC (Biolegend, clone H130,1:100, staining most hemopoietic cells), CD31-APC (Biolegend, clone WM-59, 1:100, staining endothelial cells), EPCAM-AF488 (Biolegend, clone 9C4, 1:50), CD49f-PE/Cy7 (Biolegend, clone GoH3, 1 µg ml−1, 1:200). DAPI was used to detect dead cells. Cells were filtered through a cell strainer (Partec) before sorting. Sorting of cells was done using a FACS Aria Fusion sorter. Single-stained control cells were used to perform compensation manually and unstained cells were used to set gates. After doublets, dead cells and contaminating haematopoietic and endothelial cells (referred to as lineage) were gated out (**Supplementary Fig 2**), up to 30,000 luminal progenitors were sorted for scRNAseq (with the addition of 400 HuMECs as spike-in).

### scRNA sequencing

Estimated equal numbers of cells per sample were processed for scRNA library preparation. Samples were processed for first strand cDNA synthesis after sample defrosting and single cell preparation. The remaining steps of library preparation were completed within the following 7 days.

### Library preparation and sequencing

Single-cell RNA-seq libraries were prepared in the Cancer Research UK Cambridge Institute Genomics Core Facility using the following: Chromium Single Cell 3′ Library & Gel Bead Kit v3, Chromium Chip B Kit and Chromium Single Cell 3’ Reagent Kits v3 User Guide (Manual Part CG000183 Rev C; 10X Genomics). Cell suspensions were loaded on the Chromium instrument with the expectation of collecting gel-beads emulsions containing 8,000 single cells per channel. RNA from the barcoded cells for each sample were subsequently reverse-transcribed in a C1000 Touch Thermal cycler (Bio-Rad) and all subsequent steps to generate single-cell libraries were performed according to the manufacturer’s protocol with no modifications (13 cycles were used for cDNA amplification). cDNA quality and quantity were then measured with Agilent TapeStation 4200 (High Sensitivity 5000 ScreenTape), after which 25% of material was used for gene expression library preparation.

Library quality was confirmed with Agilent TapeStation 4200 (High Sensitivity D1000 ScreenTape to evaluate library sizes) and Qubit 4.0 Fluorometer (ThermoFisher Qubit™ dsDNA HS Assay Kit to evaluate dsDNA quantity). Each sample was normalized and pooled in equal molar concentration. To confirm concentration, pools were qPCRed using KAPA Library Quantification Kit on QuantStudio 6 Flex before sequencing.

Pools were sequenced on an Illumina NovaSeq6000 sequencer with the following parameters: 28 bp, read 1; 8 bp, i7 index; and 91 bp, read 2. Four S4 flowcells were run initially (10 samples per S4 lane) and further sequencing was added (depending on number of cells captured) aiming for 50,000 reads per cell across the whole dataset.

### Processing and quality control of scRNA-seq data

The same processing and analysis pipeline was used across all samples and all batches, using both R and Python with the conda / singularity environments specified for each step in the github repository. Read processing was performed using the 10X Genomics workflow. We used the Cell Ranger Single-Cell Software Suite (v6.0.2) to perform barcode assignment, demultiplexing and UMI quantification (http://software.10xgenomics.com/single-cell/overview/welcome). The reads were aligned to the 10X reference genome GRCh38 (ref-2020-A). To demultiplex our spike-in cells we used Vireo (v0.5.6)^52^ genotyping with a reference lane of pure spike-in cells (sample SLX-20005-20446_SIGAE10), outputting the probability of being a spike-in or spike-in-doublet alongside the final classification as spike-in or not.

Barcodes that correspond to droplets with successfully captured cells were distinguished from empty droplets using the “emptyDrops” function from DropletUtils (v1.12.1)^52^ at an FDR of 0.001. Next, we performed various QC steps to identify and remove low quality cells. Starting from the filtered cellranger output we applied Scrublet (v0.2.3) in combination with an over-clustering approach by Pijuan-Sala et al. to computationally detect and remove doublets per sample^53, 54^. After an initial rough filtering on low quality cells (remove any cell with UMIs < 600 or mitochondrial content > 15%), we further excluded cells with lower than 3 MAD (Median-Absolute Deviance) from the sample median for UMIs or counts, or those above 3 MADs in mitochondrial percentage. Additionally, we found sample SLX-19864-20262_SIGAA5, belonging to Donor 50, to contain debris and have poor cell quality in general. Consequently, this sample was excluded from downstream analysis. Lastly, any spike-in cells were removed. After these filtering steps 800,198 cells were retained.

Single cell analysis was performed using the scanpy pipeline (v1.8.2)^55^. Raw counts were log-normalised (scanpy.pp.normalize_per_cell(counts_per_cell_after=1e4), scanpy.pp.log1p()) and subsequently used to select 5000 highly variable genes (scanpy.pp.highly_variable_genes()). For dimensionality reduction and batch correction the raw counts of the highly variable genes were used to train the scvi-tools (v0.17.1) scVI VAE (parameters: batch_key=“processing_date”, n_latent=20, gene_likelihood=“nb”, use_layer_norm=“both”, use_batch_norm=“none”, encode_covariates=True, dropout_rate=0.2, n_layers=2)^56, 57^. The 20 latent dimensions were used to determine a KNN graph (scanpy.pp.neighbors(n_neighbors=15)) for UMAP calculation (scanpy.tl.umap()) and leiden clustering (scanpy.tl.leiden()).

### Clustering and annotation

We used multiple rounds of iterative leiden^58^ clustering with scanpy.tl.leiden() to identify cellular partitions of our data. At each round we determined a new KNN graph and applied leiden clustering to the scVI batch corrected latent representation. This allowed us to label “level0” and “level1” annotations of the broad cellular compartments and general cell types. Then, we split our cells into multiple smaller groups, repeating the above steps to finalise the “level2” subcluster annotations. Subclusters were labelled and identified using an array of known marker genes, quality control metrics and gene lists provided by “scanpy.tl.rank_genes_groups”. We identified the optimal cluster resolution for each group maximising subcluster gene expression distinction, robustness and consistency **(Supplementary Fig 8-10**). In some cases, small clusters of additional doublets were found showing mixed lineage marker expression alongside elevated mean scrublet scores. These cells were then (re-)labelled “Doublets” at all levels of cell type annotation.

### Differential expression and gene set enrichment analysis

Differential gene expression analysis was performed using edgeR (v3.36.0)^59^. A negative binomial generalised log-linear model was fitted to gene counts with the relevant subcluster or donor groups as the test covariate(s). The subcluster marker genes (**Supplementary Fig 11a, Supplementary Table 2**) used a one-versus-all approach across all samples (10X lanes) with the following model formula: “∼ sample_type_coarse + subcluster_test” (subcluster_test being a boolean for the subcluster being tested). Tests between AR and HR-BR1/2 donors were completed separately in standard two group comparison approaches with samples again as the replicate with the following model formula: “∼ sample_type_coarse + donor_age + parity + HR_test” (HR_test being a boolean for the relevant HR donors). For each the “glmQLFTest” function was used to identify genes that have a LFC significantly different from 0 at an FDR of 0.1.

Gene set enrichment analysis based on gene ontology (GO) terms was conducted to characterise various gene sets in the analysis. We used clusterProfiler (v4.2.0) function “enrichGO” to test for GO term enrichment of the top 100 upregulated genes (ranked by log fold changes with background of all other expressed genes) defining our IFNG+ T cell population^59^.

### Milo and differential abundance analysis

To test for changes in cellular abundance at high resolution across various groups/conditions we used Milo’s (v1.3.1) neighbourhood abundance approach^24^. Due to the direct impact sorting methods have on cellular abundance we only considered unsorted (epithelial and stroma enriched) samples for all differential abundance testing. For exploring age and parity, we considered the 22 (AR) mammoplasty donors (though one was excluded due to lack of parity metadata). We generated a shared cell neighbourhoods based on the scVI corrected KNN graph for each of the epithelial, stromal and immune compartments. Specifically, we used the functions “buildGraph” and “makeNhoods” with parameters: “k=50”, “d=20”, “prop=0.3”, “refined=TRUE” and “refinement_scheme = ‘graph’” in their respective inputs. The use of a graph refinement scheme significantly decreased the run time (and removed requirement to run “calcNhoodDistance”) when testing large numbers of cells and the remaining parameters were chosen based on the recommended Milo summary plots and standards. The differential neighbourhood abundance testing was completed using “testNhoods” over samples with “fdr.weighting=’graph-overlap’” and model formula “∼ sample_type_coarse + milo_block + milo_test” with milo_block being donor_age/parous and milo_test being parous/donor_age respectively (here parous is a boolean for parity being greater than zero). To test for changes in HR donors compared to AR donors, a similar approach was taken while ensuring the neighbourhoods were consistent across both AR versus HR-BR1 and AR versus HR-BR2 tests to enable comparisons. The only differences arise in the model formula being: “∼ sample_type_coarse + parous + donor_age + milo_test” where milo_test is a boolean labelling the respective HR-BR1/2 samples. To create the ‘Milo signature’ summary we generated the average neighbourhood log fold changes per donor for each subcluster. To account for the issue of overlapping neighbourhoods, these averages were generated over the cells individually using the mean log fold change over all neighbourhoods containing the cell.

To look independently at the changes seen in our HR-unk donors we performed standard differential abundance testing using edgeR. We used a negative binomial generalised log-linear model testing cell counts per subcluster from each sample with the model formula: “∼ sample_type_coarse + parous + donor_age + HR-unk”.

### Regulon analysis

We used the python version of SCENIC^31^ with the aertslab-pyscenic-scanpy-0.12.1-1.9.1.sif singularity environment. For computational speed, we used a subset of the total epithelial populations: for each of the AR, HR-BR1 and HR-BR2 cohorts we sampled 3000 cells of each level2 subcluster (or all cells of this group if fewer than 3000 were detected). Subsequently, following the standard pipeline, we generated the gene regulatory network with “grn” then identified regulons with “ctx” and finally, the area under the curve was computed using the “aucell” function. We then used “regulon_specificity_scores” to identify the top regulons for each subcluster.

### Cell-cell interactions

To predict and analyse the putative cell-cell interactions within our dataset we used Cell Chat (v1.6.0)^41^. Here we split cells into level2 subclusters excluding the immune compartment which we grouped into lymphoid and myeloid lineages to simplify the analysis and increase readability. Separate Cell Chat objects were made for the AR, HR-BR1 and HR-BR2 cohorts to allow comparison between each. The communication probabilities were inferred using the “computeCommunProb” function. Cell–cell communications for each cell signalling pathway were generated with the “computeCommunProbPathway” function.

### Data availability

The authors declare that all data supporting the findings of this study and unprocessed images are available within the article and its supplementary information files or from the corresponding author upon reasonable request. The raw sequencing data will be made available on publication. Processed data can be found and explored using CELLxGENE at https://cellxgene.cziscience.com/collections/48259aa8-f168-4bf5-b797-af8e88da6637. All code will be made available online at GitHub.

## Supplementary Figures

## Supplementary Tables

**Supplementary Table 1 –** This table details the metadata recorded for each donor ID alongside sample (10X lane) specifics recorded during the process of sequencing. Each row represents a separate sample.

**Supplementary Table 2 –** This table lists the top 100 marker genes for each defined subcluster determined by one-vs-all pseudo-bulk testing. The top 10 genes for each subcluster are visualised as a heatmap in Supplementary Fig 11.

**Supplementary Table 3 –** This is a collection of six excel sheets detailing the differential gene expression results across our two test conditions (AR vs HR-BR1 and AR vs HR-BR2) and three epithelial cell types (LP, HS and BSL).

**Supplementary Table 4 –** This table lists the top 100 genes differentially expressed in each of the LP1-5 subclusters with respect to the remaining LP cells.

## Author contribution

ADR performed the majority of the bioinformatic analysis and interpretation of the data. SP contributed to the study design, sample processing, analysis and interpretation of the data. JS contributed to the sample processing. DK and PH contributed to the data processing, batch correction and cell cluster identification. AS contributed to the design of the sample batches and contributed to the analysis of the raw data. AJT contributed to the analysis of the data and figure design. KK performed all the scRNAseq library preparation and sequencing. RBM, IG, JJG and LJ provided the human tissues and the metadata from the 55 donors. ADR, SP, JCM and WTK wrote the paper. JCM and WTK conceptualised and supervised the study.

## Acknowledgments

This study was primarily funded by a MRC project grant (MR/S036059/1) and supported by BBSRC project grant (BB/S006745/1), Breast Cancer Now project grant (2017MayPR907), CRUK career establishment award (17348) and CRUK programme foundation award (DCRPGF\100010) to WTK and core funding from EMBL and CRUK (C9545/A29580) to JCM. We would like to thank Dr. Sandy Westermann (SCIGRAPHIX) for the illustration in Figure 1. The authors wish to acknowledge the role of the Breast Cancer Now Tissue Bank in collecting and making available the samples used in the generation of this publication, and all the patients who donated.

## Declarations

JCM has been an employee of Genentech since September 2022.

---

**Supplementary Figure 1.**
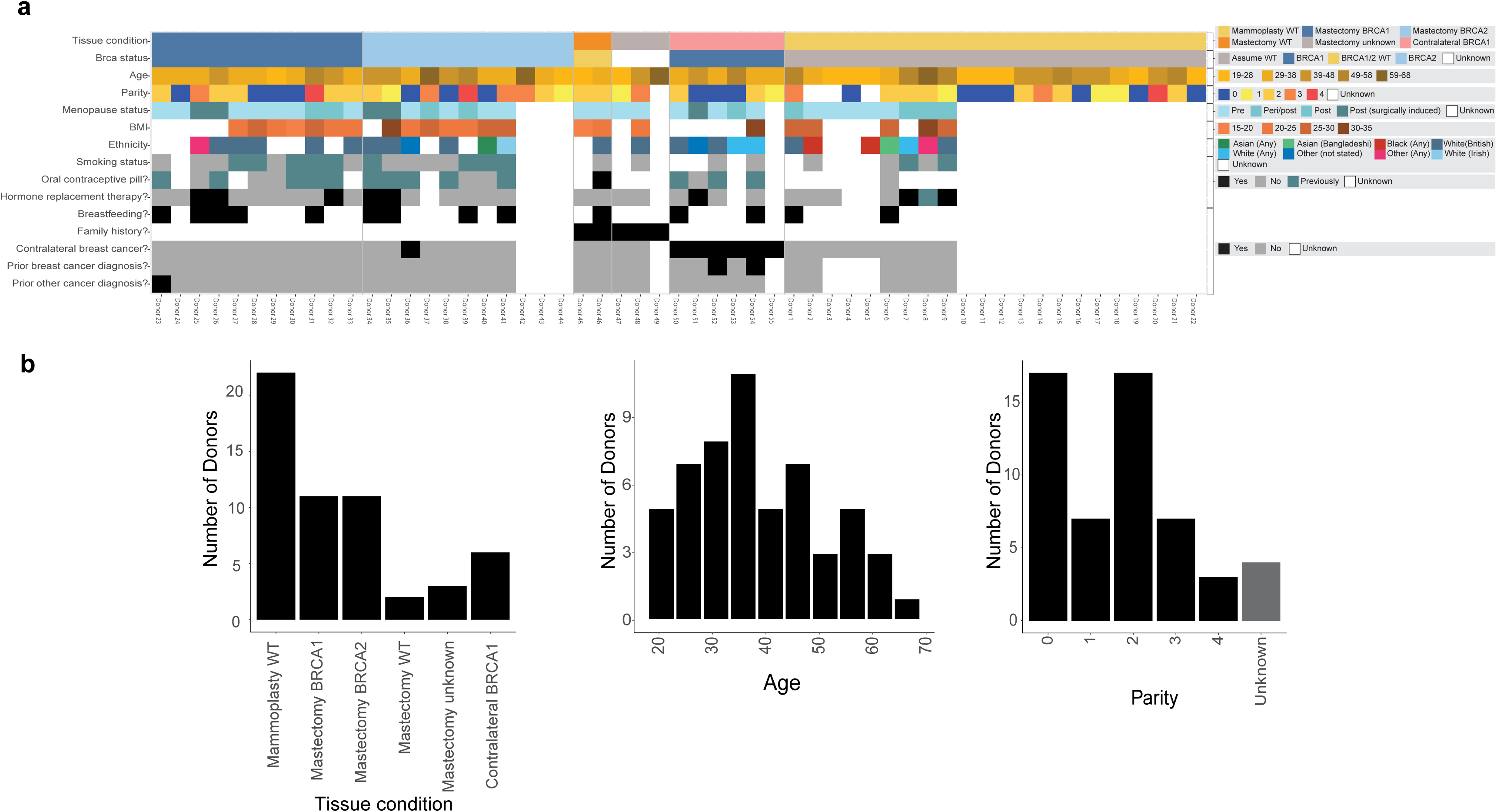
Summary of major demographic metadata. **(a)** Summary table visualising some of the main demographics metadata for each of our donors. Full metadata listed per sample (10X lane) in Supplementary Table 1. **(b)** Bar plot/histograms showing the total distribution of our main three demographics explored in the paper – tissue condition (surgery and BRCA status), age and parity.

**Supplementary Figure 2.**
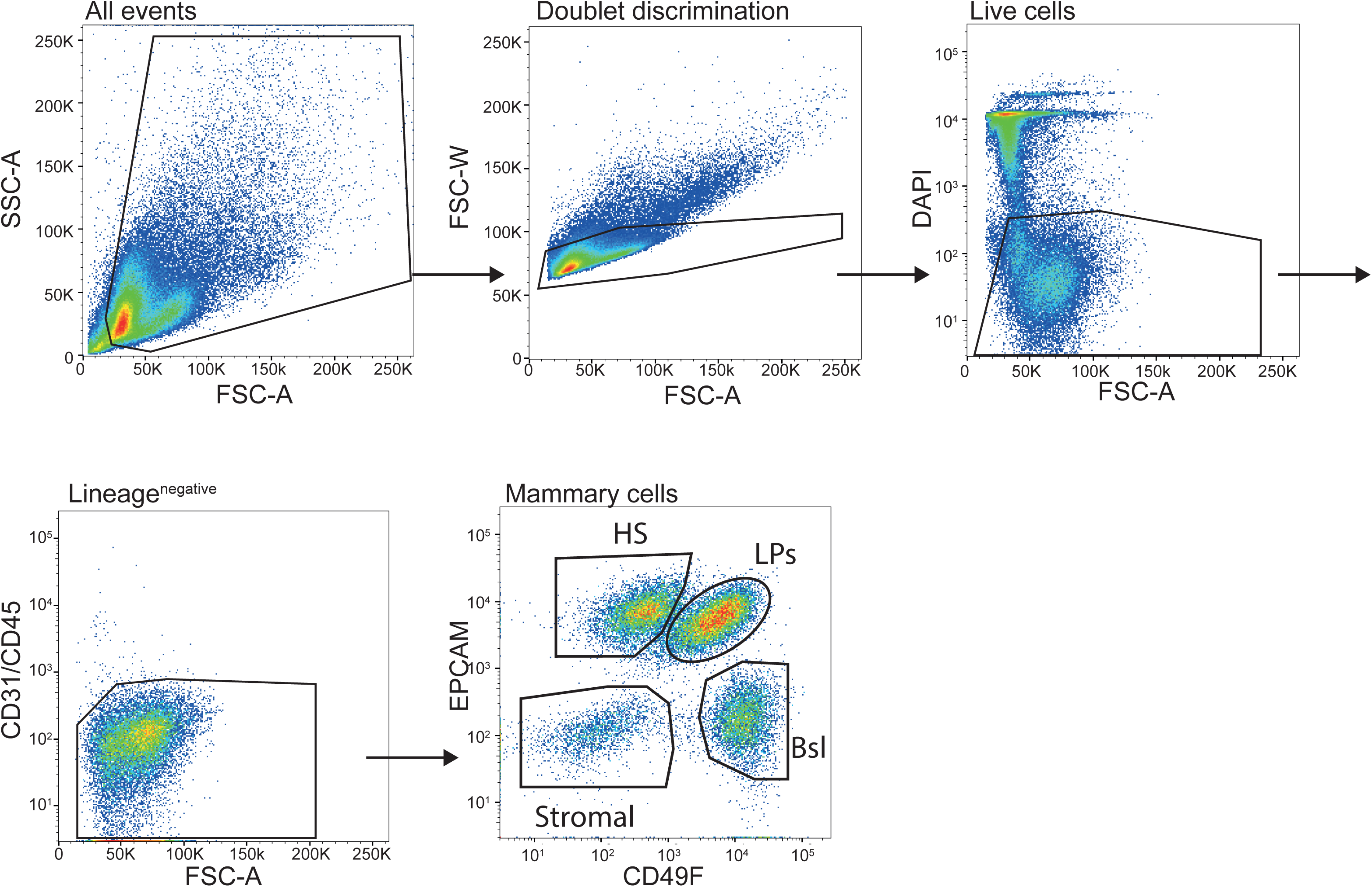
Gating strategy used for human samples on the FACS Aria Fusion sorter. Representative plots showing the gating strategy used to select live, lineage negative, single luminal progenitor cells based on EPCAM and CD49f staining of single cell preparations. FSC-W: forward scatter width, FSC-A: forward scatter area, SSC-A: side scatter area. The arrows indicate sequential gating. This gating strategy was used for all samples.

**Supplementary Figure 3.**
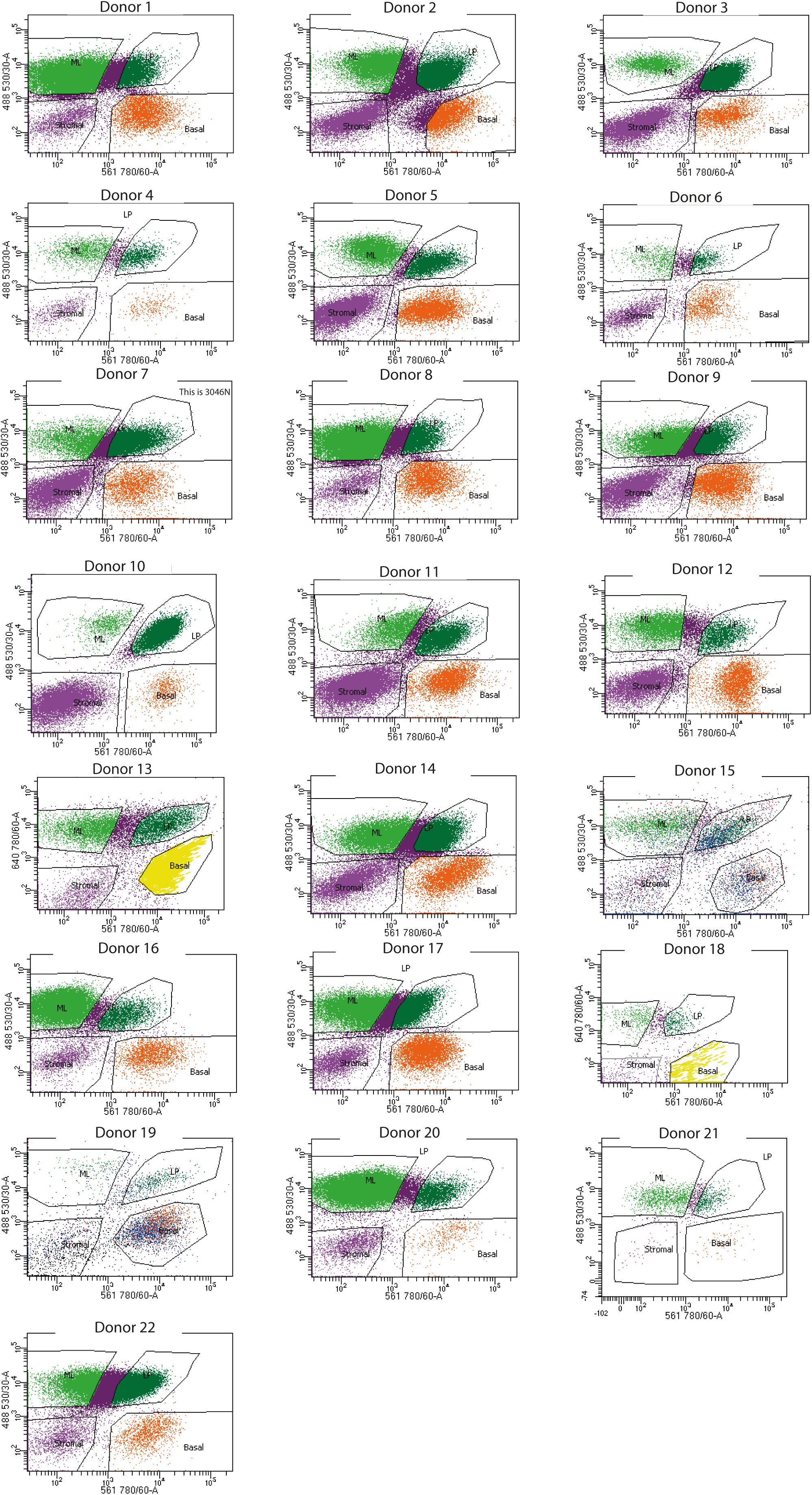
Individual samples FACS plots for mammoplasty donors. FACS plots showing EPCAM (shown as 488 530/30-A) and CD49f (shown as 561 780/60-A) expression on live, lineage negative, single cells for individual mammoplasties samples. ML indicates hormone sensing (HS) population. Donor IDs are indicated on each plot.

**Supplementary Figure 4.**
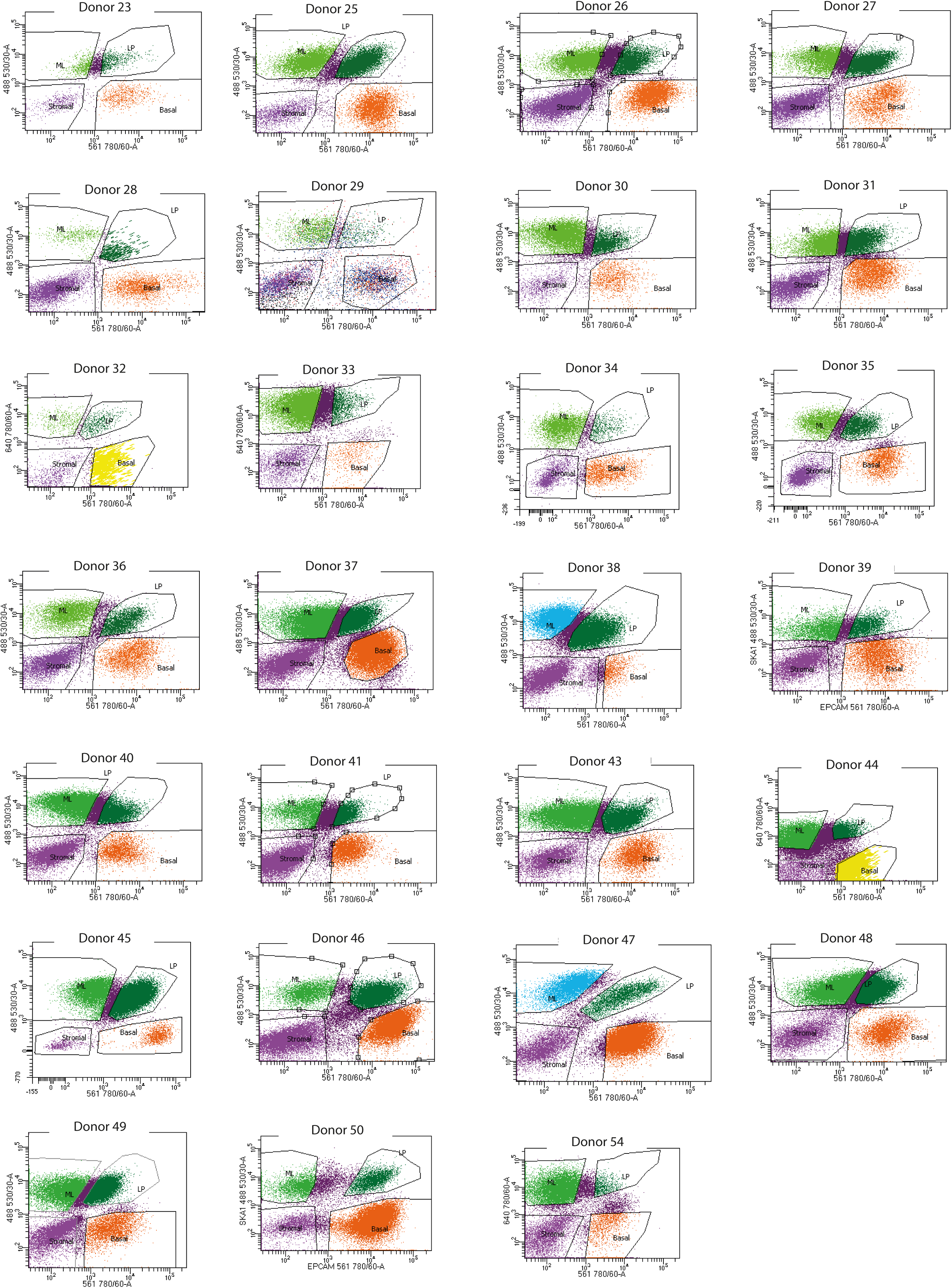
Individual samples FACS plots for mastectomy and contralateral donors. FACS plots showing EPCAM (shown as 488 530/30-A) and CD49f (shown as 561 780/60-A) expression on live, lineage negative, single cells for individual mastectomies and contralateral samples. ML indicates hormone sensing (HS) population. Donor IDs are indicated on each plot.

**Supplementary Figure 5.**
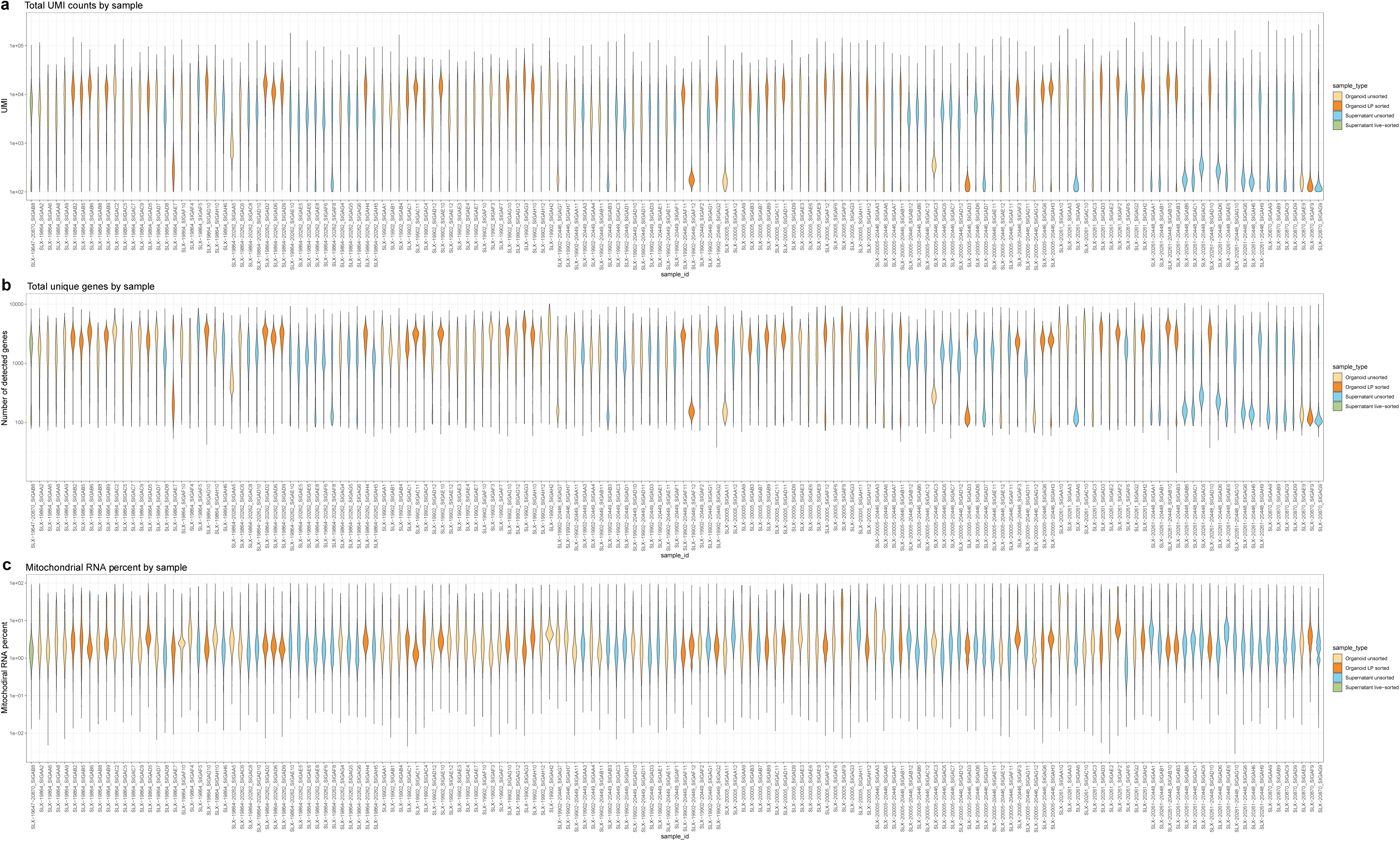
Pre-filter sample quality control metrics. **(a)** A collection of violin plots for the quality control unique molecular identifiers (UMI) counts per 10X lane (sample_id) prior to the relevant cutoffs (methods). **(b)** A collection of violin plots for the quality control (unique) gene counts per 10X lane (sample_id) prior to the relevant cutoffs (methods). **(c)** A collection of violin plots for the quality control mitochondrial RNA proportions per 10X lane (sample_id) prior to the relevant cutoffs (methods).

**Supplementary Figure 6.**
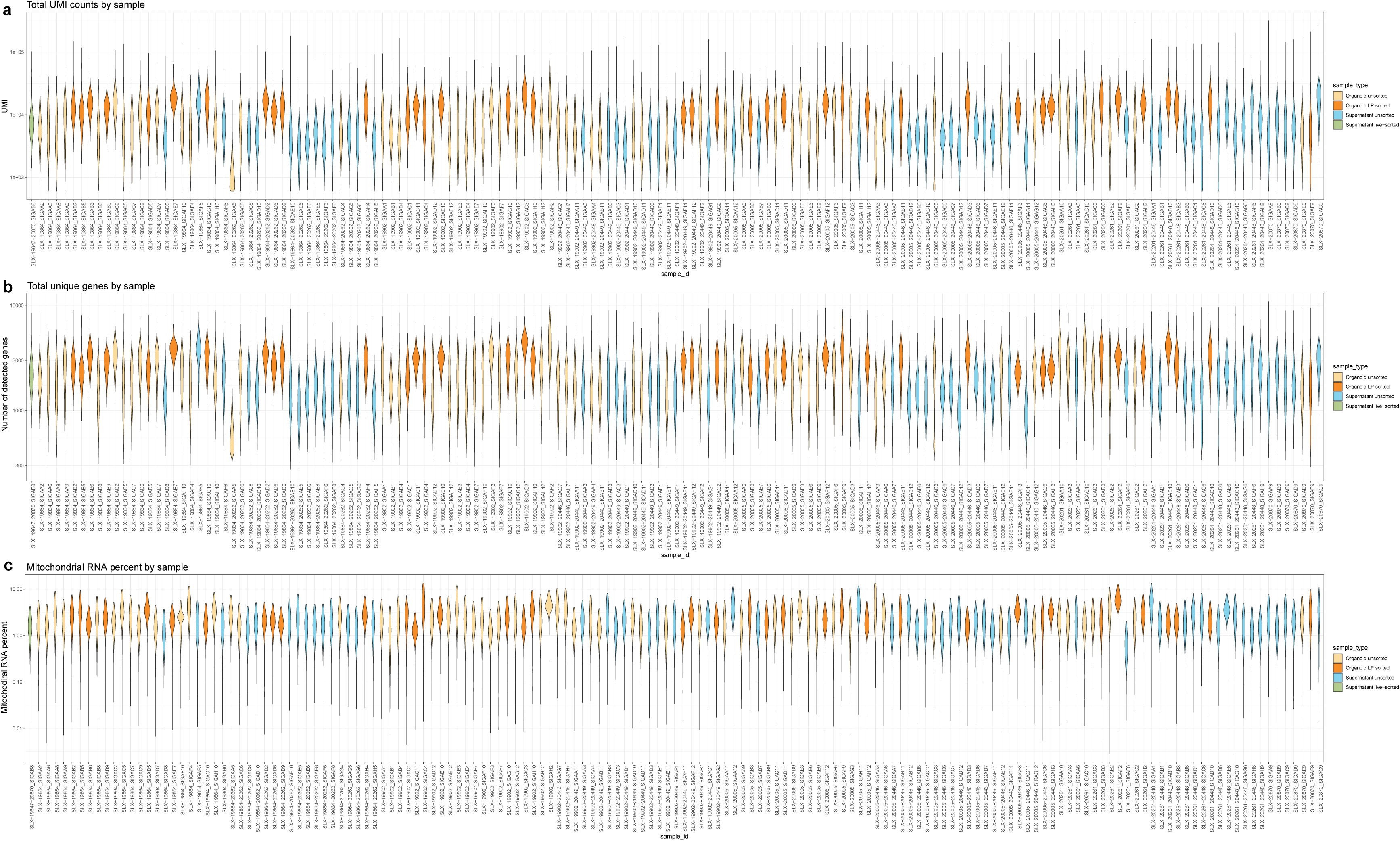
Post-filter sample quality control metrics. Same as **Supplementary Figure 5** but for the cleaned (post-filter) cells passing the necessary quality control thresholds (see methods).

**Supplementary Figure 7.**
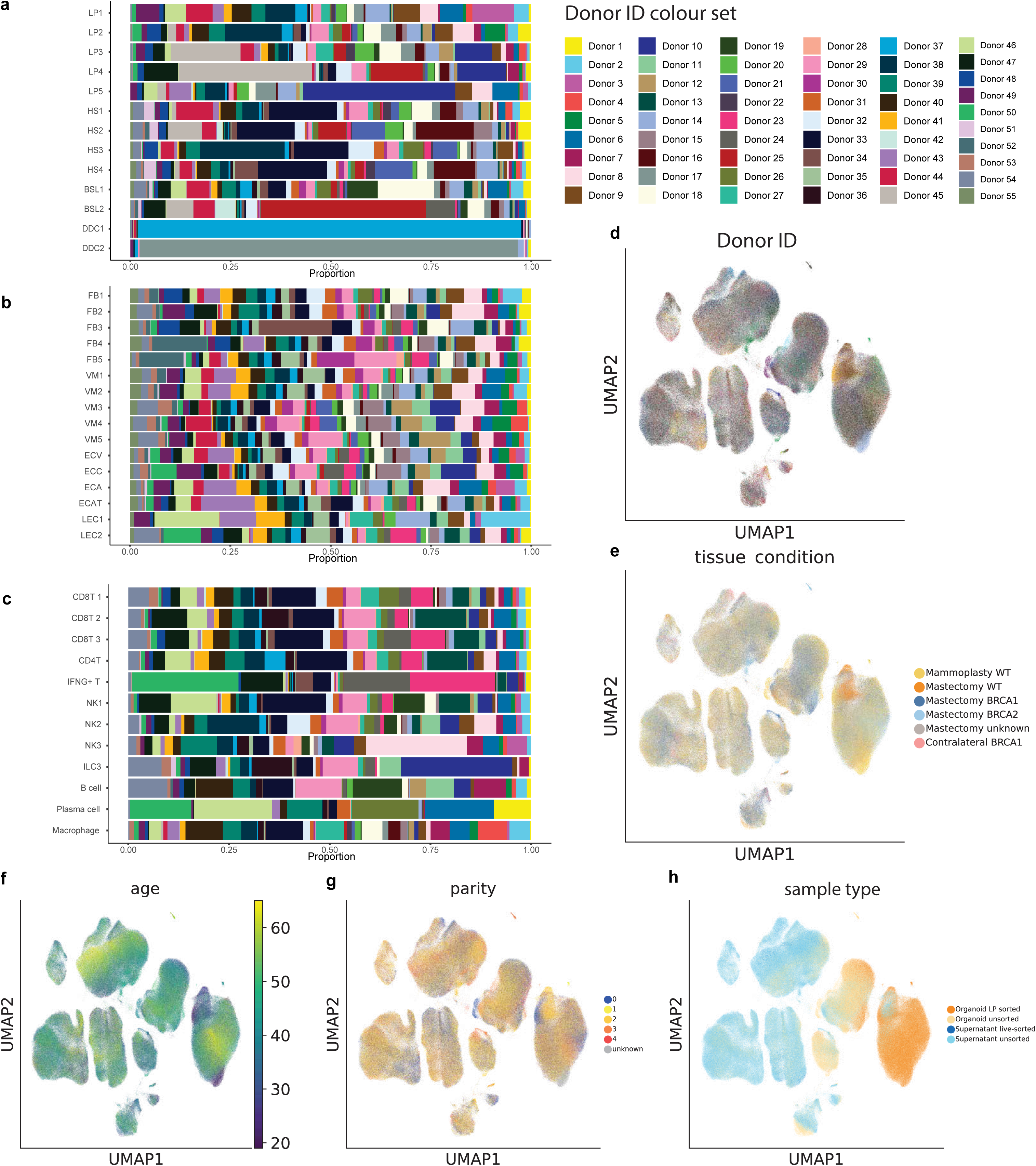
Subcluster donor and demographic heterogeneity. **(a-c)** Bar plots displaying the proportions of donor contribution per cell type subcluster for the epithelial (top), stromal (middle) and immune (bottom) compartments respectively. **(d-h)** Uniform manifold approximation and projection (UMAP) plots of all cells coloured by their respective donor, tissue condition (collection surgery performed and BRCA status), age, parity and sample type (see Figure 1 for types of ‘enriched cell’ sample types).

**Supplementary Figure 8.**
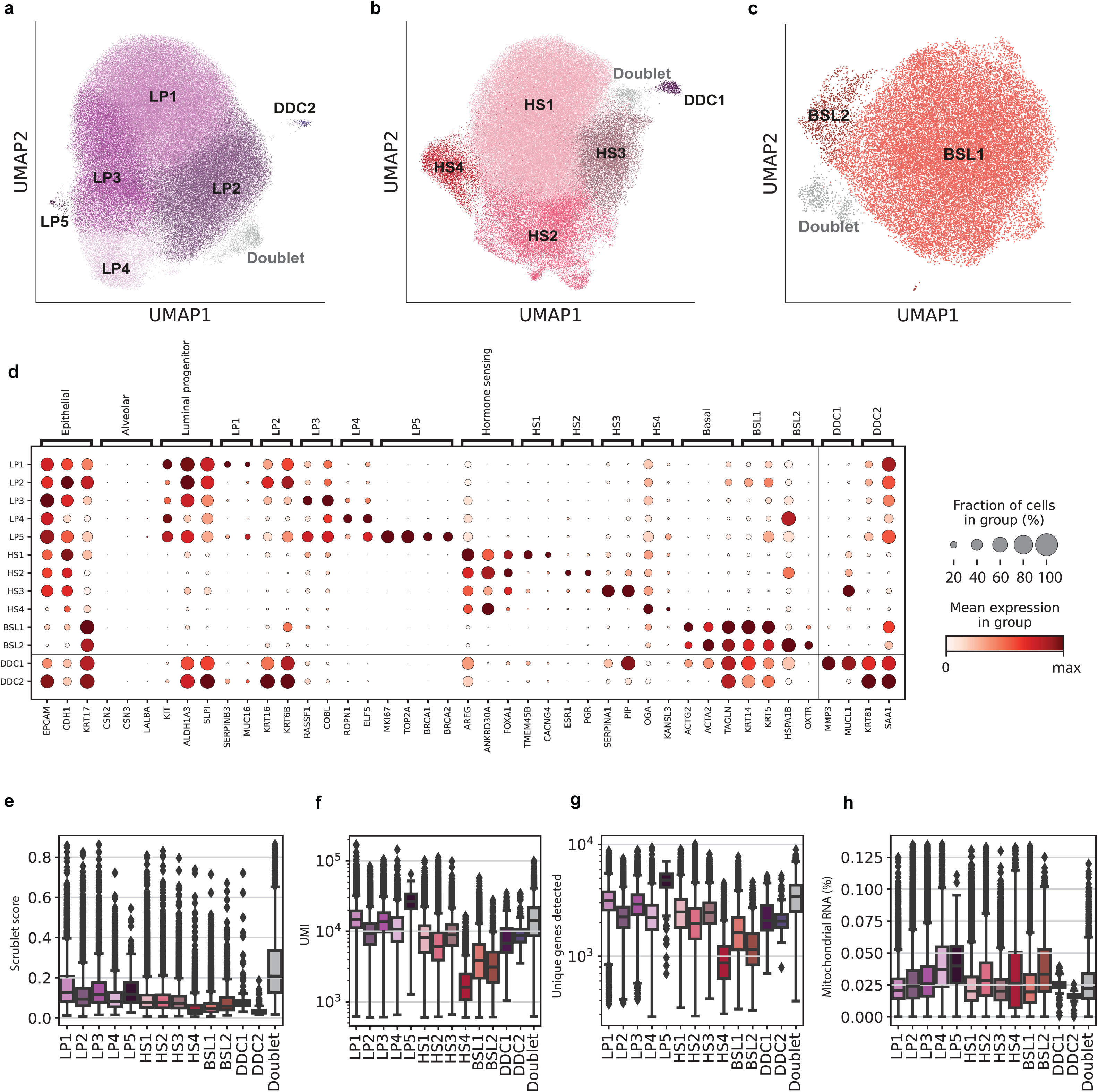
Epithelial cell type and subcluster annotation. **(a-c)** Cell type specific uniform manifold approximation and projections (UMAPs) for the luminal progenitor, hormone sensing and basal cell types used to identify subclusters. **(d)** A dot plot summarising the known markers used to identify cell types and selection of genes distinguishing each subcluster for the epithelial compartment. Each row corresponds to a specific cell subcluster and its expression of each gene (column) normalised per gene, brackets at the top detail the cell type/subcluster that this gene marks. **(e-h)** Boxplots showing the distribution of Scrublet (doublet) scores, unique molecular identifier (UMI) counts, unique gene counts and mitochondrial RNA percentages across the epithelial subclusters. Combined with marker gene expression these were used to identify any additional doublet clusters annotated in (a-c).

**Supplementary Figure 9.**
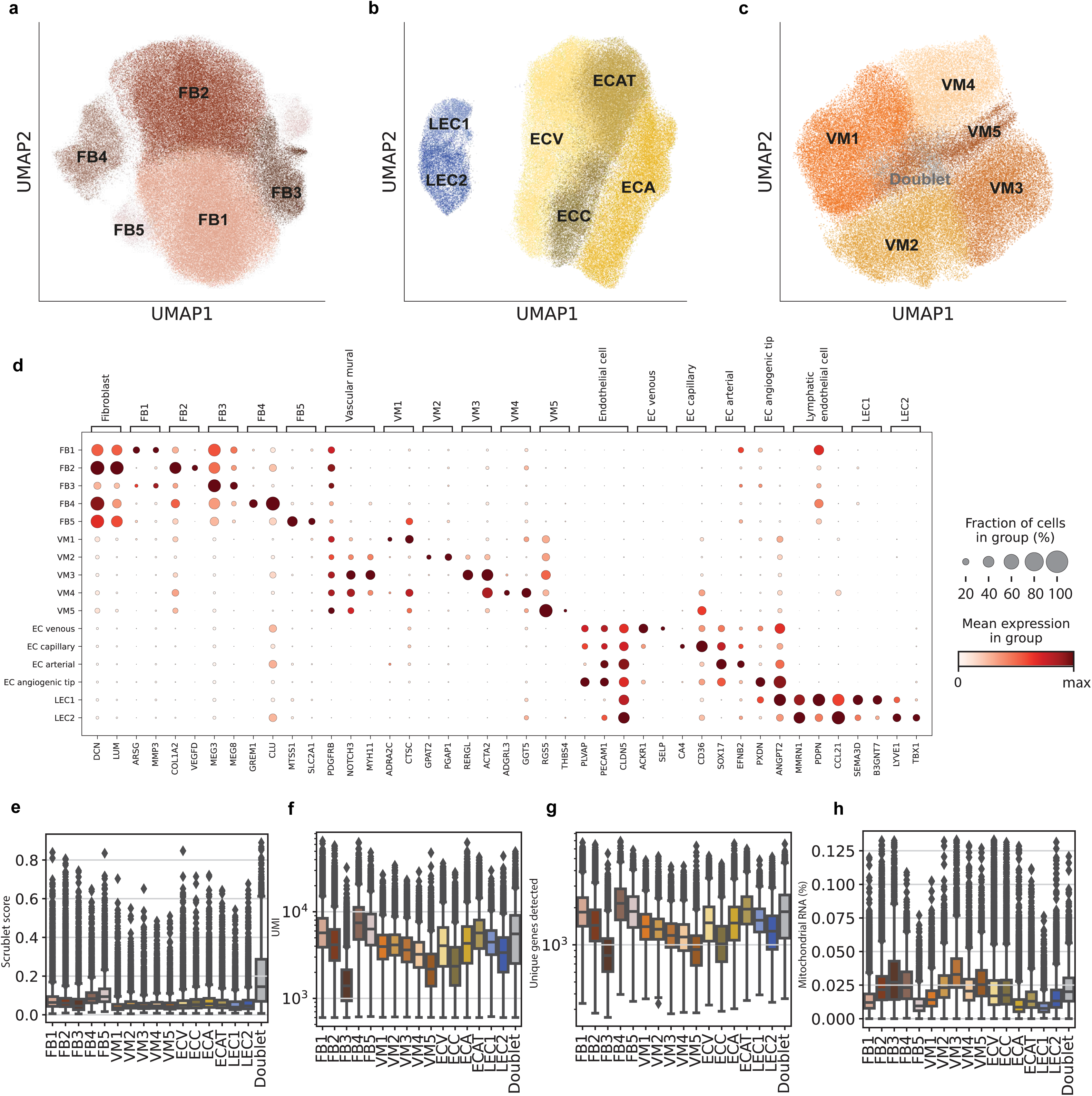
Stromal cell type and subcluster annotation. **(a-c)** Cell type specific uniform manifold approximation and projections (UMAPs) for the fibroblast, endothelial and vascular mural cell types used to identify subclusters. **(d)** A dot plot summarising the known markers used to identify cell types and a selection of genes distinguishing each subcluster for the stromal compartment. Each row corresponds to a specific cell subcluster and its expression of each gene (column) normalised per gene, brackets at the top detail the cell type/subcluster that this gene marks **(e-h)** Boxplots showing the distribution of Scrublet (doublet) scores, unique molecular identifier (UMI) counts, unique gene counts and mitochondrial RNA percentages across the stromal subclusters. Combined with marker gene expression, these were used to identify additional doublet clusters annotated in (a-c).

**Supplementary Figure 10.**
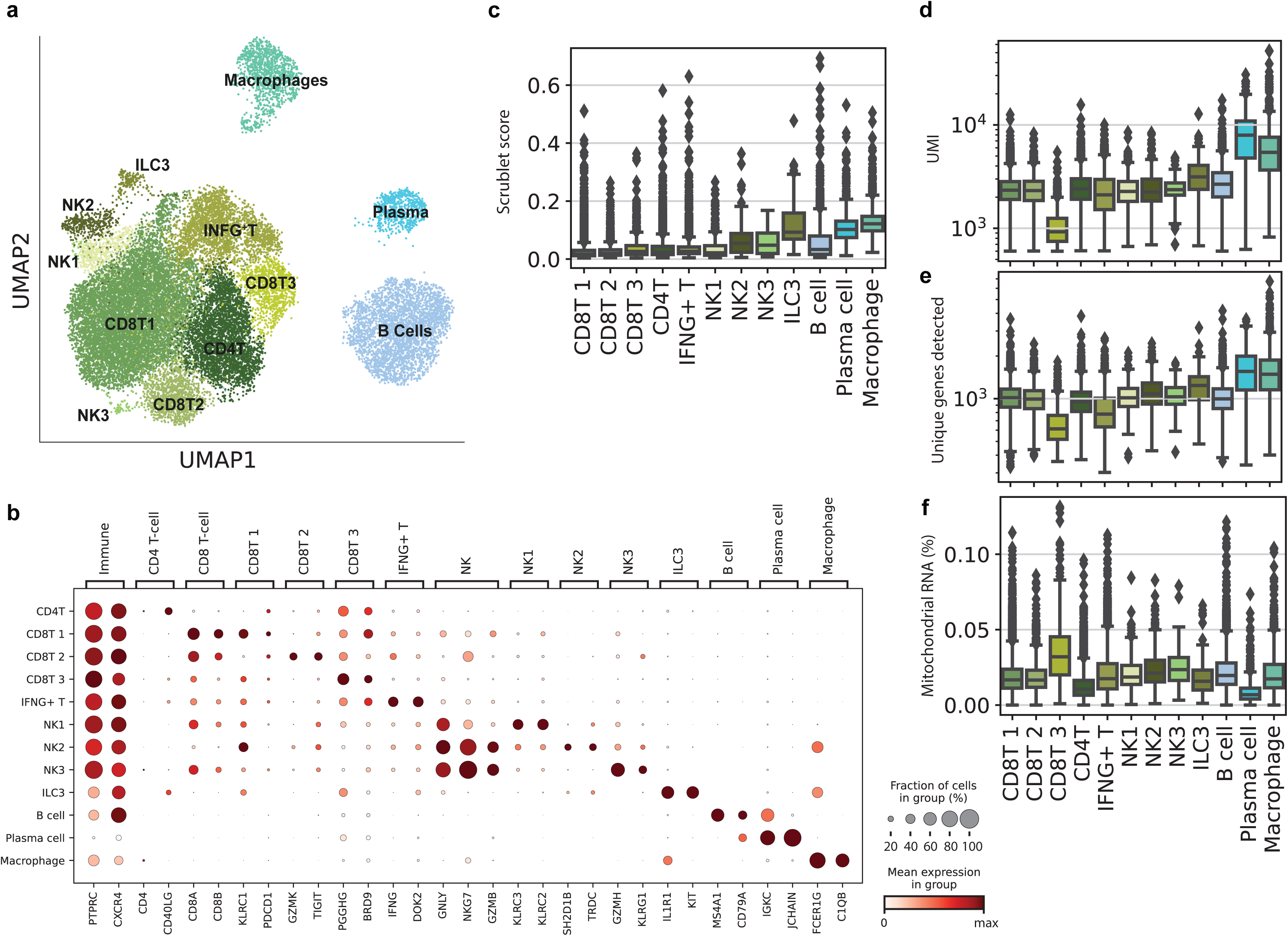
Immune cell type and subcluster annotation. **(a)** Cell type specific uniform manifold approximation and projections (UMAPs) for the immune cell types used to identify subclusters. **(b)** A dot plot summarising the known markers used to identify cell types and a selection of genes distinguishing each subcluster for the immune compartment. Each row corresponds to a specific cell subcluster and its expression of each gene (column) normalised per gene, brackets at the top detail the celltype/subcluster that this gene marks. (**c-f)** Boxplots showing the distribution of Scrublet (doublet) scores, unique molecular identifier (UMI) counts, unique gene counts and mitochondrial RNA percentages across the immune cell subclusters. Unlike the epithelial and stromal compartments, these show no doublet clusters identified here and that they were removed in earlier stages.

**Supplementary Figure 11.**
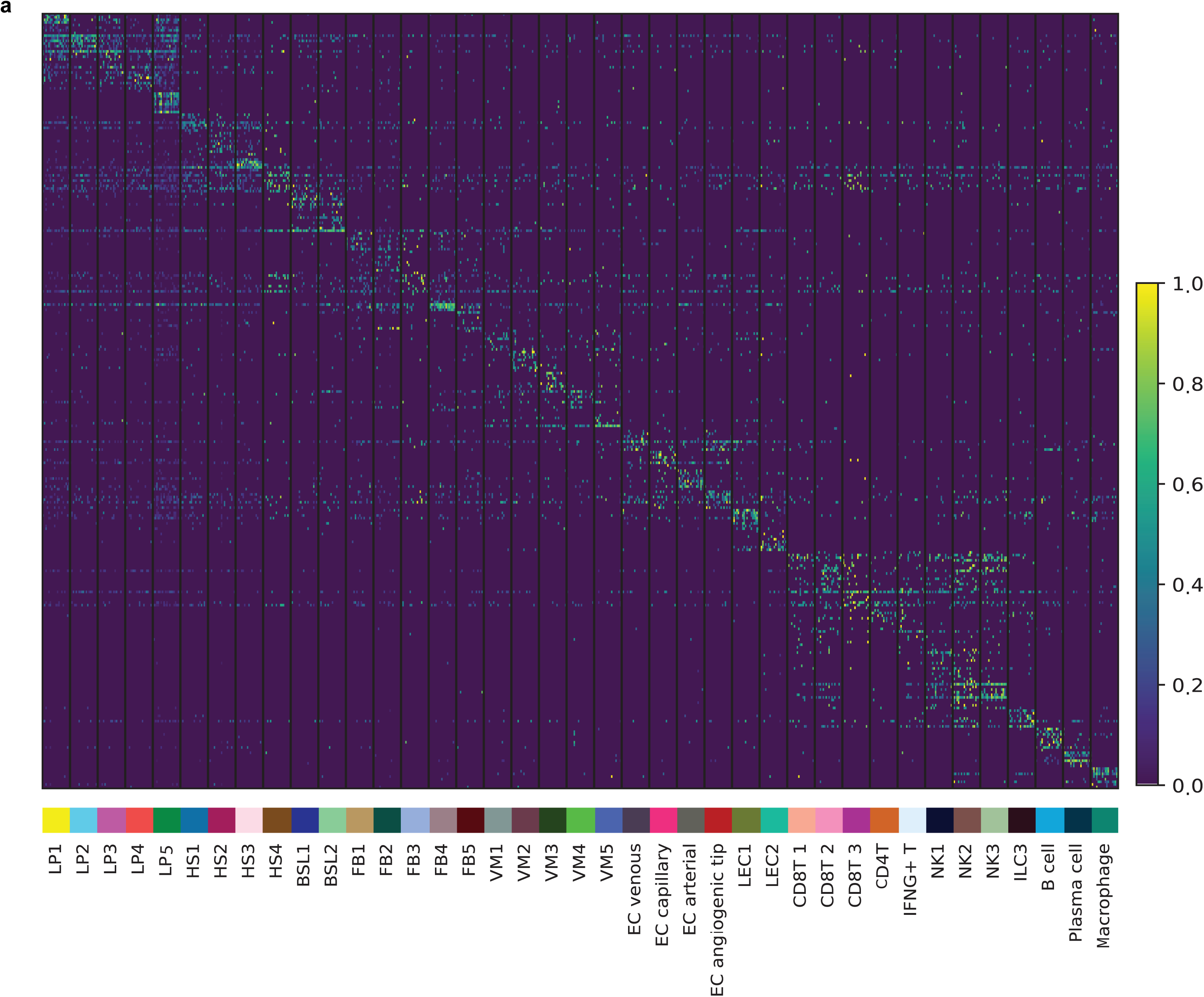
Specificity of subcluster gene signatures. **(a)** Heatmap showing the top 10 marker genes (rows) for each subcluster identified via several one vs all negative binomial tests for each subcluster (columns; subsetted to 100 cells per subcluster). Full lists of the top 100 genes per subcluster are detailed in **Supplementary Table 2**. The heatmap was made using scanpy.pl.heatmap() with parameters “standard_scale=’var’” and “layer=’logcounts’”.

**Supplementary Figure 12.**
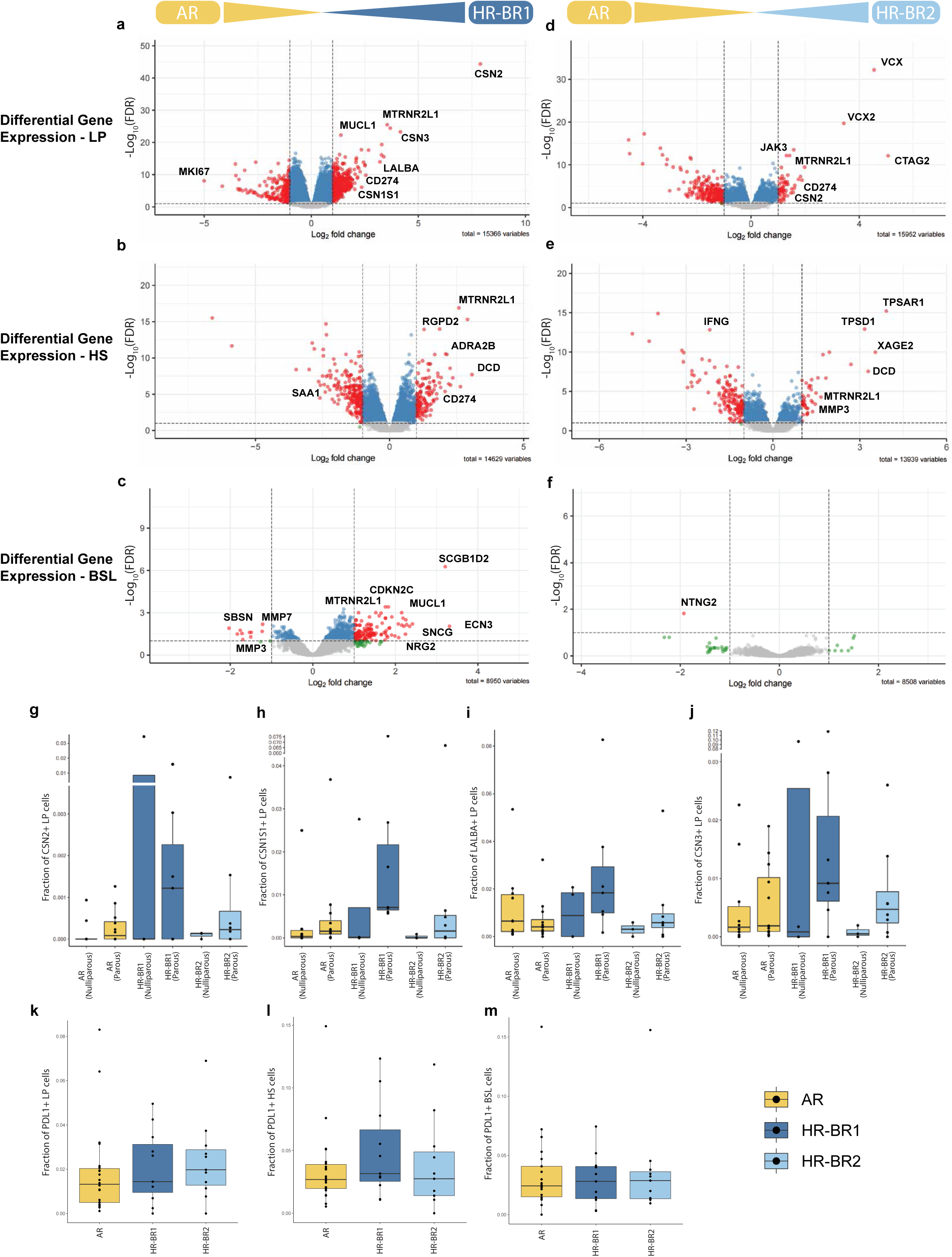
Differential gene expression of high-risk donor epithelium. **(a-c)** Volcano plots showing the results of differential gene expression testing between average risk (AR) and high risk BRCA1 germline (HR-BR1) donors within the luminal progenitor (LP), hormone sensing (HS) and basal (BSL) compartments respectively. Green points have significant log fold changes, blue points have significant FDR (FDR < 0.1), red points have significant log fold change and FDR. **(d-f)** Volcano plots showing the results of differential gene expression testing between AR and high risk BRCA2 germline (HR-BR2) donors within the luminal progenitor, hormone sensing and basal compartments respectively. Green points have significant log fold changes, blue points have significant FDR (FDR < 0.1), red points have significant log fold change and FDR. **(g-j)** Box plots showing the upregulation of milk biosynthesis related genes in HR donors compared to AR alongside the related impact of parity on the expression of these genes. These plots show the frequency of non-zero expression of each respective gene amongst the LP population. **(k-m)** Box plots showing the upregulation of PDL1 expression in HR donors compared to AR in the LP, HS and BSL cell types. These plots show the frequency of non-zero expression of PDL1 in each cell type.

**Supplementary Figure 13.**
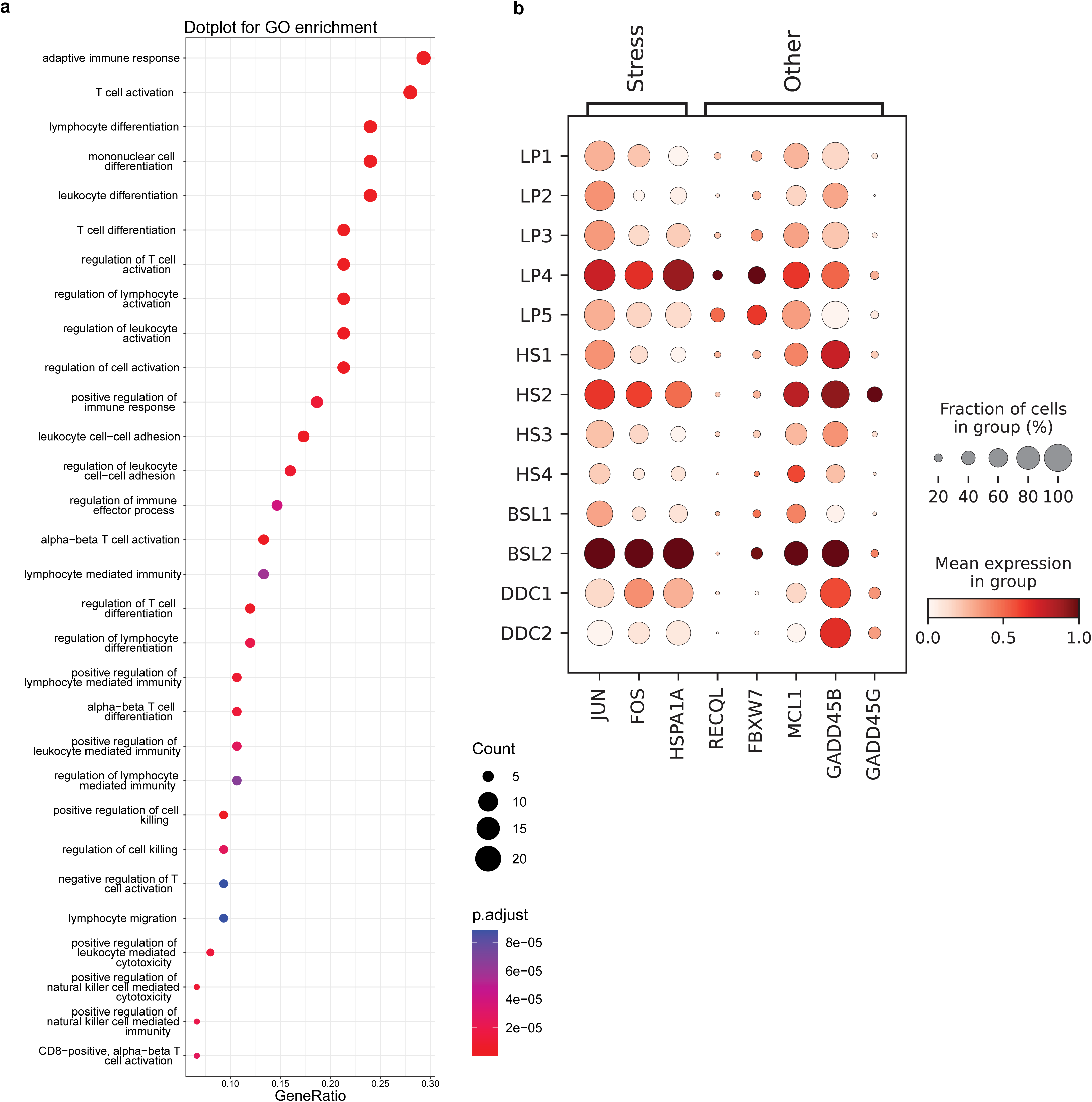
INFG+ T cell GO enrichment and LP4/HS2/BSL2 additional markers. **(a)** Gene ontology (GO) enrichment results (see methods) for the top 100 genes distinguishing the IFNG+ T cell from Supplementary Table 2. **(b)** A dot plot showing a selection of additional markers shared between LP4, HS2 and BSL2. Each row corresponds to a specific cell subcluster and its expression of each gene (column), brackets at the top detail the cell type/subcluster that this gene marks.

**Supplementary Figure 14.**
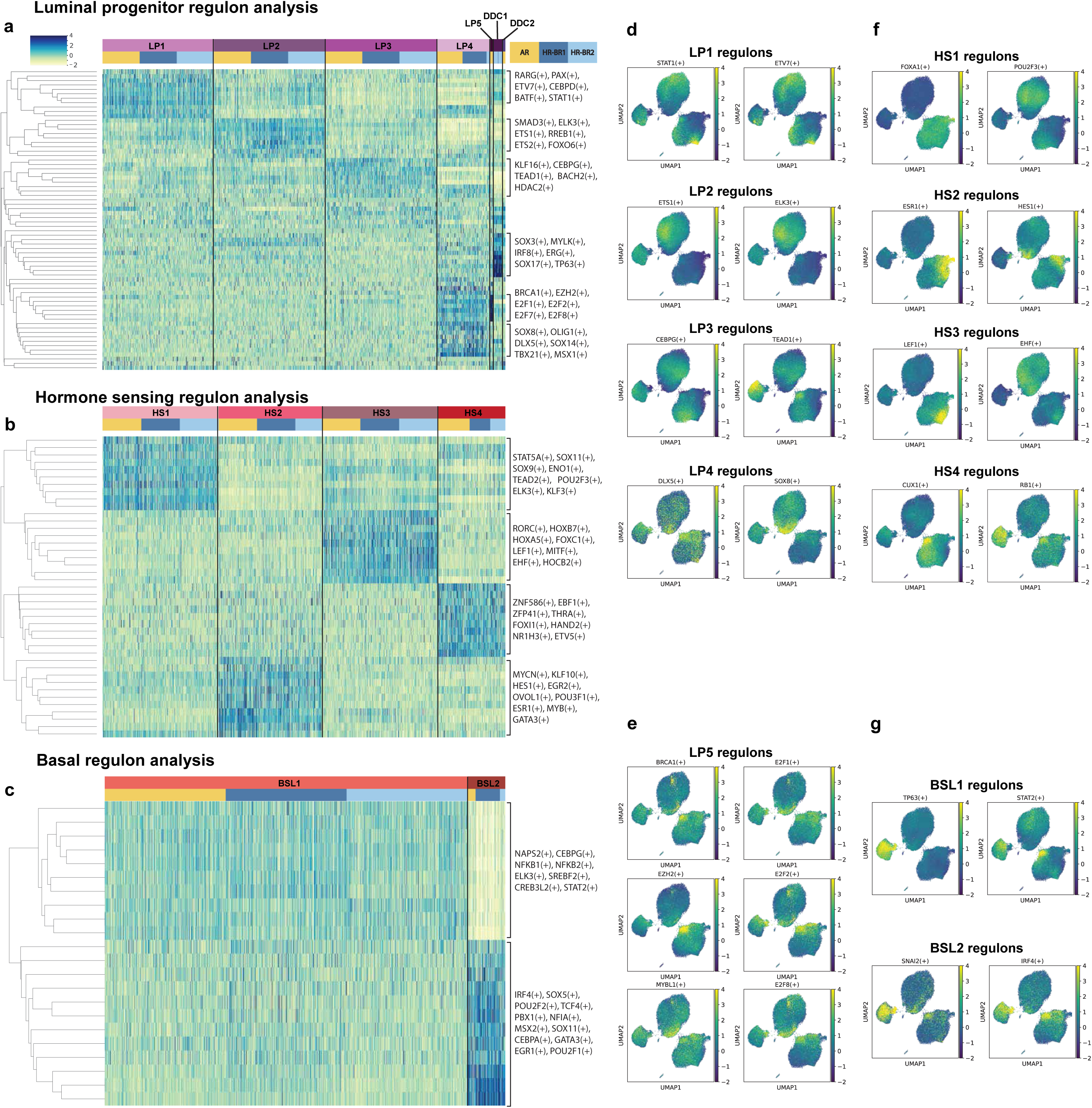
SCENIC regulon analysis of the epithelium. **(a-c)** A selection of heatmaps displaying the major regulons identified by SCENIC that are regulating the gene expression of each specified epithelial subcluster within the luminal progenitor, hormone sensing and basal cell types. The cells are first ordered by level2 subclusters then by risk condition between average risk (AR), high risk BRCA1 germline (HR-BR1) and high risk BRCA2 germline (HR-BR2) cohorts. **(d)** A collection of uniform manifold approximation and projections (UMAPs) displaying the distribution of a selection of the major regulons identified by SCENIC for the LP1-4 subclusters. **(e)** A collection of UMAPs displaying the distribution of a selection of the major regulons identified by SCENIC for the LP5 subcluster. **(f)** A collection of UMAPs displaying the distribution of a selection of the major regulons identified by SCENIC for the HS1-4 subclusters. **(g)** A collection of UMAPs displaying the distribution of a selection of the major regulons identified by SCENIC for the BSL1-2 subclusters.

**Supplementary Figure 15.**
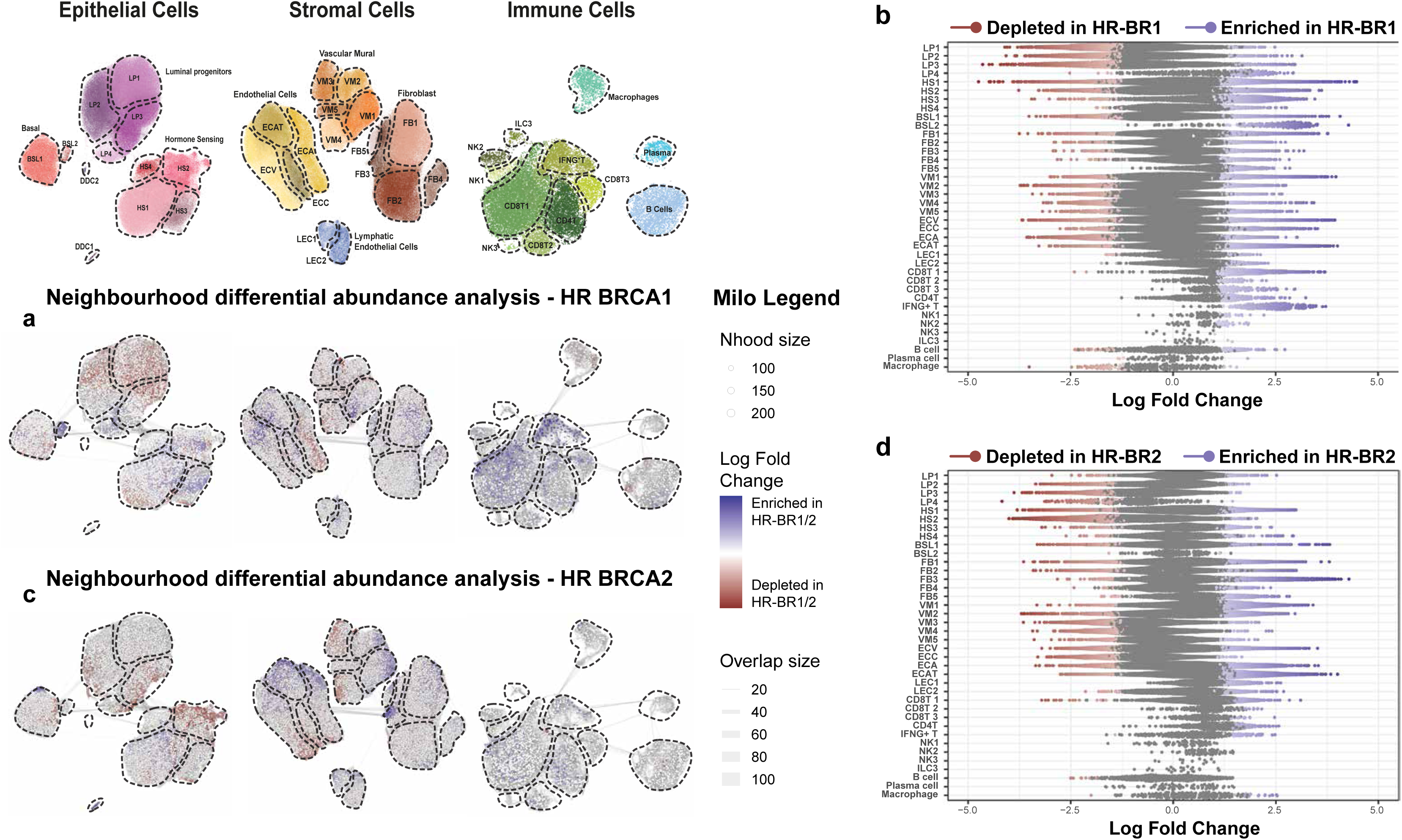
Impact of BRCA1 and BRCA2 germline mutations on the cellular composition of the breast. **(a)** Milo cell neighbourhood differential abundance plots of the significant (FDR < 0.05) changes in the breast composition comparing average risk (AR) donors (n=22) to the high risk BRCA1 germline (HR-BR1) cohort (n=11), blocking for the effects of age and parity. Blue represents enrichment with HR-BR1 whilst red denotes depletion with HR-BR1. **(b)** Beeswarm plot of the log fold changes of the Milo neighbourhoods grouped into each cell type subcluster for AR versus HR-BR1. Neighbourhoods with a significant change in cellular abundance are coloured as indicated. **(c)** Milo cell neighbourhood differential abundance plots of the significant (FDR < 0.05) changes in the breast composition comparing AR donors (n=22) to the high risk BRCA2 germline (HR-BR2) cohort (n=11), blocking for the effects of age and parity. Blue represents enrichment with HR-BR2 whilst red denotes depletion with HR-BR2. **(d)** Beeswarm plot of the log fold changes of the Milo neighbourhoods grouped into each cell type subcluster for AR versus HR-BR2. Neighbourhoods with a significant change in cellular abundance are coloured as indicated.

**Supplementary Figure 16.**
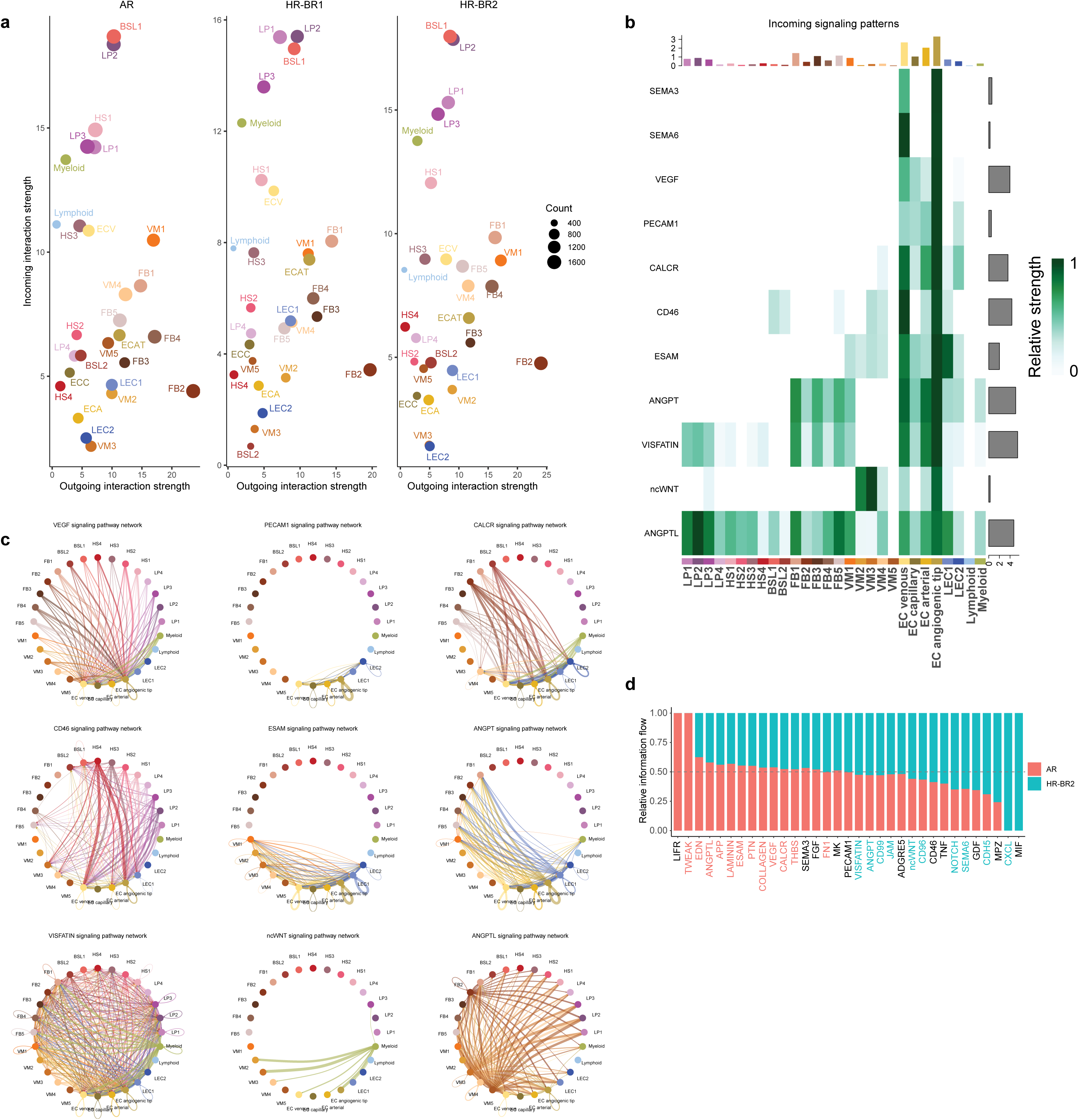
Cell-cell communication of high-risk donors. **(a)** Scatter plot showing the overall changes in total incoming and outgoing cell-cell interactions (through Cell Chat predicted ligand-receptor interactions) comparing average risk (AR) to high risk BRCA1 germline (HR-BR1) and high risk BRCA2 germline (HR-BR2) donors. **(b)** A heatmap displaying the 12 interaction pathways with most specific incoming signal to the endothelial angiogenic tip cells. **(c)** Circle plots of 10 of the top 12 most specific incoming signalling pathways to endothelial tip cells (the top two are plotted in Figure 4d). The lines indicating directed cell-cell communication are coloured by the ‘sending’ cell subcluster and lead towards the receiving cell subcluster and their width is determined by the number of these predicted interactions. **(d)** A relative information flow bar plot showing the relative changes in proportions of cell-cell communication pathways in AR and HR-BR2 donors. The pathway axis labels are coloured blue/red if they are significantly (p < 0.05) up/downregulated in HR-BR2 donors.

**Supplementary Figure 17.**
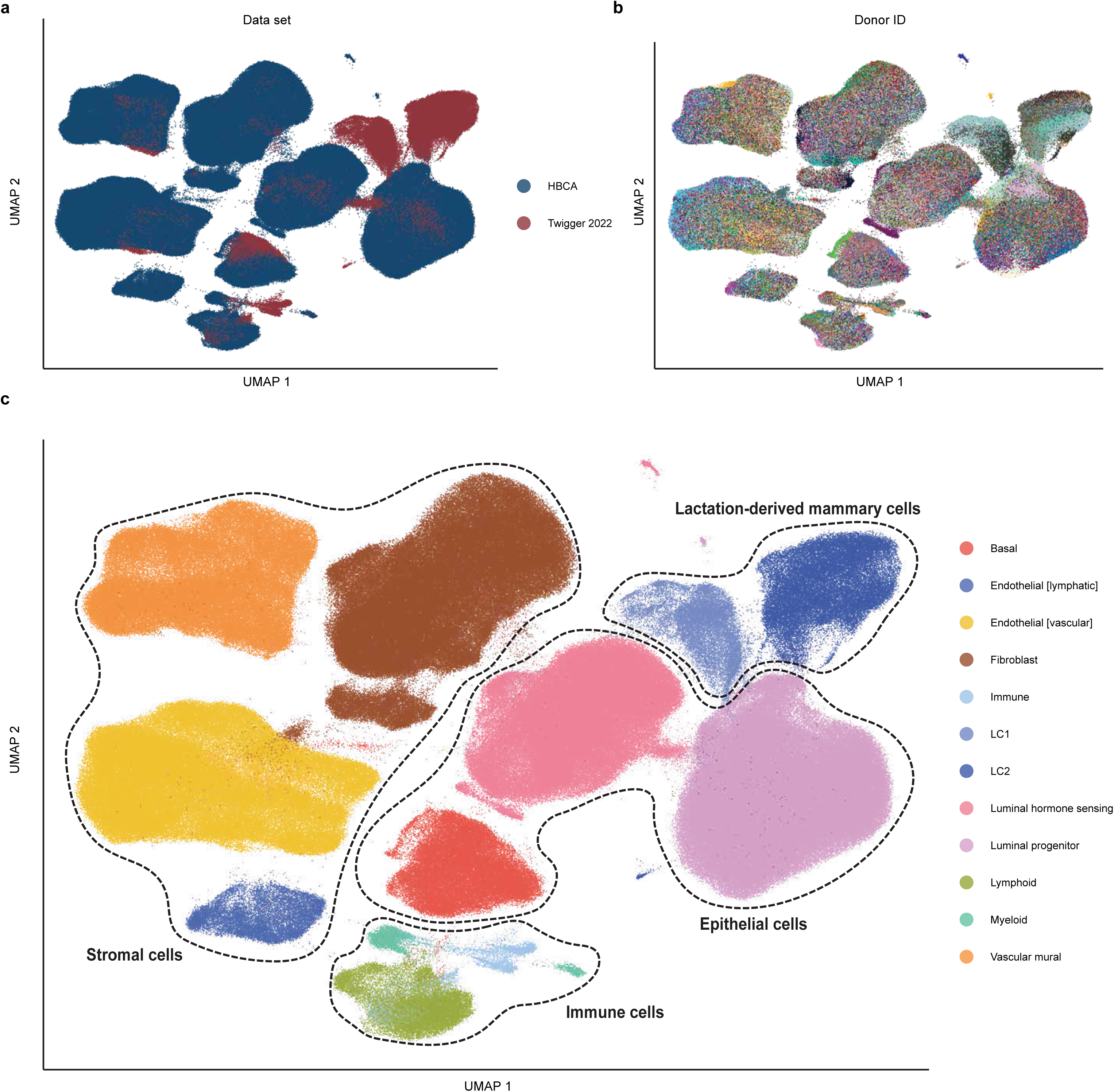
Atlas growth capability and integration to include lactation-derived mammary cells. **(a-c)** Uniform manifold approximation and projection (UMAP) plots of the batch integrated HBCA and Twigger 2022 data sets coloured by the dataset of origin, donor ID and combined cell type annotation respectively.

